# Loci under balancing selection facilitate the emergence of pseudo-overdominance and recombination suppression

**DOI:** 10.1101/2025.06.19.660549

**Authors:** Lou Guyot, Tatiana Giraud, Paul Jay

## Abstract

Loci under strong balancing selection, such as sex-determining, mating-type, and self-incompatibility loci, are frequently flanked by regions of suppressed recombination. The reasons why recombination suppression evolves around these loci remain poorly understood. Here, we propose that one reason may be that loci under balancing selection facilitate the emergence of pseudo-overdominance, a phenomenon under which linked partially recessive deleterious mutations in repulsion mimic overdominance. Pseudo-overdominance arises under certain conditions when linkage disequilibrium gradually builds up between recessive deleterious mutations, often due to strong genetic linkage and genetic drift, forming complementary haplotypes. Homozygous and recombinant offspring are then selected against because they carry homozygous recessive deleterious mutations. Using individual-based simulations, we demonstrate here that the presence of loci under balancing selection (overdominant) eases the establishment of pseudo-overdominance by facilitating the maintenance of partially recessive deleterious mutations and linkage disequilibrium in their flanking regions. This occurs particularly in regions with low recombination rates and a high load of partially recessive deleterious mutations, which can be found in specific genomic regions in natural populations. We additionally show that selfing, while generally hindering pseudo-overdominance establishment, can in contrast facilitate its establishment around a permanently heterozygous locus, likely because recessive deleterious mutations are protected from being purged near such a locus and selfing increases genetic drift. Once established, we further showed that pseudo-overdominance can in turn favor the evolution of recombination suppression around permanently heterozygous loci, preventing the creation of unfit homozygous recombinant offspring. These results suggest new avenues for understanding the evolution of loci under balancing selection, sex chromosomes, mating-type chromosomes in selfing fungi, supergenes, and shed new light on the mechanisms driving genome evolution.

## Introduction

Recombination is fundamental in evolution as it generates new allelic combinations upon which selection can act. Yet, recombination is suppressed in many genomic regions, either entirely or between some haplotypes (Schwander et al. 2014; Dapper and Payseur 2017; Stapley et al. 2017; Peñalba and Wolf 2020). Textbook examples include sex chromosomes, mating-type chromosomes, self-incompatibility loci, centromeric regions, as well as many supergenes (B. Charlesworth 1991; D. Charlesworth 2016, 2017; Schwander et al. 2014; Fernandes et al. 2019; Hartmann, Duhamel, et al. 2021; Vittorelli et al. 2023). Beyond these well-known cases, many other genomic regions display heterogeneous recombination landscapes, with localized hotspots interspersed among regions of low recombination rates (Jeffreys et al. 2001), as observed in the major histocompatibility complex in mammals (Yoo et al. 2025). While such genomic regions with low recombination rates can show low diversity due to hitchhiking effects (e.g. in pericentromeric regions; B. Charlesworth and Jensen 2021), they are often associated with some form of balancing selection, maintaining high levels of genetic diversity (Kamau and D. Charlesworth 2005; Küpper et al. 2016; Carey et al. 2021; Hartmann, Duhamel, et al. 2021; Jay et al. 2021; Le Veve et al. 2023).

The evolutionary drivers of recombination suppression, and the extent to which they are lineage-or context-dependent, remain long-standing questions in evolutionary biology (Wright et al. 2016; Ponnikas et al. 2018; Jay et al. 2024). For sex chromosomes, it has long been considered that stepwise extension of recombination suppression evolves because it allows linkage between sex-determining loci and sexually antagonistic loci, i.e. loci with alleles beneficial to one sex but deleterious to the other, or even with mere different selection intensities between sexes (D. Charlesworth and B. Charlesworth 1980; Rice 1987; Bergero and D. Charlesworth 2009; D. Charlesworth 2016, 2017; Flintham et al. 2025). A similar, more general scenario can be formulated for many supergenes and other loci under balancing selection, where recombination suppression would evolve to preserve beneficial combinations of co-adapted alleles at epistatic loci (Fisher 1930; Berdan et al. 2023) or even at non-epistatic loci under selection for local adaptation across the same contrasting environments (Kirkpatrick and Barton 2006; B. Charlesworth and Barton 2018). Because of its simplicity and intuitive adaptive rationale, this model is appealing. Yet, decades of research on sex chromosomes have provided limited empirical support for the sexual antagonism hypothesis (Beukeboom et al. 2014; Wright et al. 2016, 2017; Ponnikas et al. 2018; Jay et al. 2024; Ironside 2010). Signals of antagonistic selection may however be hard to detect in genomes. Powerful methods have recently been developed (Connallon and Clark 2012, 2014), allowing for example to detect signals of sexually antagonistic variation in *Drosophila* sex chromosomes, although without any enrichment compared to autosomes (Ruzicka et al. 2025).

More importantly, the sexual antagonism hypothesis cannot account for the progressive recombination suppression around fungal mating-type loci. The initial establishment of recombination suppression in fungi can be explained in basidiomycete fungi by the advantage of linking mating-type loci and/or mating-type loci and the centromere, as it increases gamete compatibility odds in selfing species with two mating-type loci (Branco et al. 2017; Hartmann, Duhamel, et al. 2021). However, this is not the case in ascomycete fungi that carry a single mating-type locus. For example, recombination suppression has evolved independently around the mating-type locus in at least four distinct lineages of Sordariales fungi, without involving linkage to either a centromere or another mating-type locus (Hartmann, Duhamel, et al. 2021; De Filippo et al. 2026). Furthermore, recombination suppression often extends stepwise away from mating-type loci (Branco et al. 2017; Hartmann, Duhamel, et al. 2021; De Filippo et al. 2026; Lucotte et al. 2025). The sexual antagonism hypothesis cannot explain this extension beyond mating-type loci and centromeres, because i) no form of antagonistic selection is known between mating types: no particular trait is known, or can be imagined, to be more beneficial in one mating type compared to the alternative one, beyond mating-type functions *per se* (Branco et al. 2017; Bazzicalupo et al. 2019; Hartmann, Duhamel, et al. 2021), and ii) mating types govern mating compatibility at the haploid stage in fungi, and all the fungi in which recombination suppression has been reported beyond mating-type loci are precisely those without any significant haploid phase in nature, so that there is very little room for antagonistic selection between different mating types in the life cycle (Branco et al. 2017; Hartmann, Duhamel, et al. 2021; Jay et al. 2024; De Filippo et al. 2026). For example, in the self-fertile Sordariales fungi (*Podospora, Neurospora tetrasperma,* and *Schizothecium* fungi), nuclei of opposite mating types are packaged together into the same ascospore immediately after meiosis, during ascospore formation. As a result, the meiotic products (ascospores) are already dikaryotic, each containing two nuclei of opposite mating types, and the organism develops directly as a dikaryon (see, for example, Fig. 3 in Hartmann, Duhamel, et al. 2021). The fungi with a long haploid phase, of alternative mating types, in which there could possibly be room for antagonistic selection between mating types, in contrast show no recombination suppression around their mating-type locus (Jay et al. 2024; De Filippo et al. 2026). This highlights the need to consider alternative mechanisms that could drive the evolution of recombination suppression on sex chromosomes and other genomic regions (Jeffries et al. 2021; Lenormand and Roze 2022; Jay et al. 2024, 2025).

Here, we propose that recombination suppression can evolve around some loci under balancing selection because this form of selection could promote the emergence of a phenomenon known as pseudo-overdominance in nearby regions. Pseudo-overdominance refers to the apparent overdominance of a locus or genomic region that arises, under certain conditions, not from genuine heterozygous advantage at any particular site, but from linkage disequilibrium between multiple sites with partially recessive deleterious mutations arranged in repulsion; this configuration creates complementary haplotypes, e.g., one haplotype with + − + − and the other with − + − +, with + being wild-type alleles and – recessive deleterious mutations (B. Charlesworth and D. Charlesworth 1997; Pamilo and Pálsson 1998; Zhao and B. Charlesworth 2016; Gilbert et al. 2020; Waller 2021; B. Charlesworth 2022; Abu-Awad and Waller 2023). Homozygotes for these haplotypes experience a strong fitness disadvantage while heterozygotes have their load sheltered to some extent. As a result, heterozygotes are favoured and each site with a deleterious mutation behaves as if it was overdominant, so that genetic diversity is maintained at high levels within these “pseudo-overdominant zones”. Although recombination is not necessarily truly suppressed in regions of pseudo-overdominance, recombinant haplotypes incur a fitness cost. This is because individuals with recombinant haplotypes tend to be homozygous for numerous deleterious mutations, due to the disruption of haplotype complementation. As a result, the effective recombination rate is *de facto* reduced in these regions, not by true recombination suppression, but by strong selection against recombinant haplotypes. This might eventually select for mechanisms truly suppressing recombination, such as chromosomal inversions, that would prevent the formation of unfit progeny.

The occurrence of pseudo-overdominance depends on the buildup of linkage disequilibrium between partially recessive deleterious mutations typically driven by drift and genetic linkage inducing Hill-Robertson interferences (Hill and Robertson 1966; Ohta and Kimura 1969; Felsenstein 1974; Gilbert et al. 2020; B. Charlesworth and Jensen 2021; Waller 2021; Abu-Awad and Waller 2023). Theoretical works have shown that pseudo-overdominance is, as a result, more likely to arise in regions of low recombination rates, in species with small effective population sizes, under high mutation rates and when deleterious mutations are strongly recessive (Ohta and Kimura 1969; Gilbert et al. 2020; Waller 2021; Abu-Awad and Waller 2023). While the existence of pseudo-overdominant zones essentially remains a theoretical prediction, empirical studies in natural populations have recently identified genomic regions characterized by low recombination rates, high genetic diversity and the presence of recessive deleterious mutations, that may thus be shaped by pseudo-overdominance. Notable examples include centromeric regions (Altemose et al. 2022), the MHC locus in humans (van Oosterhout 2009), the region surrounding the mating-type locus in *Sordariales* and *Microbotryum* fungi (Menkis et al. 2008; Hartmann, Ament-Velásquez, et al. 2021; Hartmann, Duhamel, et al. 2021; Guyot et al. 2025; Branco et al. 2018; Hood and Antonovics 2000; Thomas et al. 2003; De Filippo et al. 2026), the supergene controlling wing colour patterns in the butterfly *Heliconius numata* (Jay et al. 2021), as well as large regions of the pearl millet genome (Salson et al. 2025), bird genomes (Ishigohoka et al. 2024) and the human genome (Gilbert et al. 2020).

Loci under balancing selection have been proposed to promote the emergence of pseudo-overdominance zones (Abu-Awad and Waller 2023). By maintaining distinct haplotypic backgrounds, balancing selection effectively partitions the population into groups with reduced effective population sizes, allowing different neutral or slightly deleterious mutations to accumulate in linked regions. Consistent with this, loci under balancing selection have been shown to promote the maintenance of linked deleterious mutations at intermediate frequencies (Leach et al. 1986; Antonovics and Abrams 2004; Uyenoyama 2005; Llaurens et al. 2009; Lenz et al. 2016; Glemin 2022; Tezenas et al. 2023) . Although this does not constitute pseudo-overdominance in itself, it may facilitate the evolution of pseudo-overdominance zones, i.e. the formation of complementary haplotypes carrying different sets of deleterious mutations in repulsion. However, the conditions under which balancing selection favours the establishment of pseudo-overdominance remain unexplored.

It also remains unknown how selfing might influence pseudo-overdominance establishment, despite its relevance to understand recombination suppression around mating-type loci in fungi, as those with non-recombining mating-type chromosomes are often highly selfing (Branco et al. 2017; Hartmann, Ament-Velásquez, et al. 2021; Jay et al. 2024; De Filippo et al. 2026). While selfing facilitates the purge of recessive deleterious mutations (Lande and Schemske 1985; Glémin 2003), theoretical work has shown that such mutations can be maintained for long near a permanently heterozygous locus, even under selfing (Tezenas et al. 2023). In addition, selfing also strengthens genetic drift by reducing the effective population size (Pollak 1987; Glémin 2007), which could promote pseudo-overdominance establishment.

In addition, it remains to investigate whether pseudo-overdominance, once established, can select for true recombination suppression, in particular around loci under balancing selection. Indeed, recombination is deleterious under pseudo-overdominance, as it produces homozygous unfit offspring, so that mechanisms truly suppressing recombination might be selected for.

Here, aiming at filling these gaps, we used multi-locus, individual-based simulations to investigate how, why and under what conditions pseudo-overdominant zones can emerge around loci under balancing selection. Given the wide variation across the tree of life, and even among regions within genomes, in terms of gene density, gene structure, mutation rate, recombination rate, distribution of fitness effects and dominance coefficients of mutations, and given the substantial uncertainty in their estimates, we did not aim to model population-specific parameter values. Instead, we sought to obtain a general understanding of the conditions favouring pseudo-overdominance establishment. We investigate in particular the effect of selfing and modelled scenarios reflecting mating-type loci, sex-determining loci and other overdominant loci. We first demonstrate that the sojourn time and heterozygosity of partially recessive deleterious mutations is higher in their vicinity, even under selfing. We then show that this maintenance of recessive deleterious mutations around overdominant loci expands the range of parameter values allowing the formation of pseudo-overdominant zones. While selfing generally impairs pseudo-overdominance establishment, it could favor it to some extent around permanently heterozygous loci. Finally, we show that, because pseudo-overdominance leads to reduced fitness of recombinant offspring, it can favour the evolution of true recombination suppression on mating-type chromosomes and related architectures. We further discuss that the conditions favoring the emergence of pseudo-overdominance (e.g., small population sizes and low recombination rates) might be met in particular genomic regions in certain species.

## Results

Aiming at determining whether and how loci under balancing selection promote the establishment of pseudo-overdominance, we began by focusing on diploid organisms with a bi-allelic, permanently heterozygous locus, subjected to an extreme form of overdominant selection, representative of randomly mating diploid species with two mating types, such that only gametes of opposite mating types can fuse (such as in some fungi). We also investigate the effect of selfing and other population genetic parameters on the establishment of pseudo-overdominance, in the presence or the absence of such a permanently heterozygous locus. We then extend our analysis to other types of loci under balancing selection, i.e. to a sex-determining locus mimicking proto-XY systems or proto-ZW systems, and other overdominant loci with various strengths of selection. We used simulations under a Wright-Fisher model, i.e. with a constant and finite population size, as well as discrete and non-overlapping generations, in which parents are sampled each generation with probabilities proportional to their fitness.

### Permanently heterozygous loci facilitate the maintenance of heterozygous recessive deleterious mutations in their flanking regions

Because pseudo-overdominance relies on the presence of linked recessive deleterious mutations, we first explored in detail how the presence of bi-allelic permanently heterozygous loci affected the dynamics of such mutations in their flanking regions. Using individual-based simulations, we introduced a single mutation at varying genetic distances from a permanently heterozygous locus in a population of 1000 diploid individuals, the selfing rate being initially set to 0.

As expected following previous studies (Glémin et al. 2001; Llaurens et al. 2009; Lenz et al. 2016; Tezenas et al. 2023; Hudson 1990; Zeng et al. 2021), we found that mutations persisted longer when they were closer to the permanently heterozygous locus (Fig 1A). For example, a partially recessive deleterious mutation with selection and dominance coefficients of *s*=-0.005 and *h*=0.1 was maintained about 50 times longer when at 1 × 10⁻⁶ cM from the permanently heterozygous locus than when totally unlinked to it, i.e., at 50 cM. We also found that mutations were maintained more frequently in the heterozygous state when they were closer to the permanently heterozygous locus (Fig 1B). For example, a partially recessive deleterious mutation with selection and dominance coefficients of *s*=-0.005 and *h*=0.1 exhibited an approximately 7-fold greater excess of heterozygosity when at 1 × 10⁻⁶ cM from the permanently heterozygous locus than when totally unlinked to it. Additionally, as expected, the less deleterious and the more recessive a mutation was, the longer it persisted and remained in the heterozygous state. The slight excess of observed over expected heterozygosity at unlinked loci likely results from the different sampling assumptions underlying these two measures, whereas the small decrease in mean sojourn time at very high linkage may partly result from the finite duration and stochastic variability of the simulations (see Supplementary Text 1). The dynamics of deleterious mutations unraveled here around permanently heterozygous loci should facilitate the build-up of a cluster of multiple recessive deleterious alleles in repulsion and thus promote the establishment of pseudo-overdominance.

**Figure 1:**
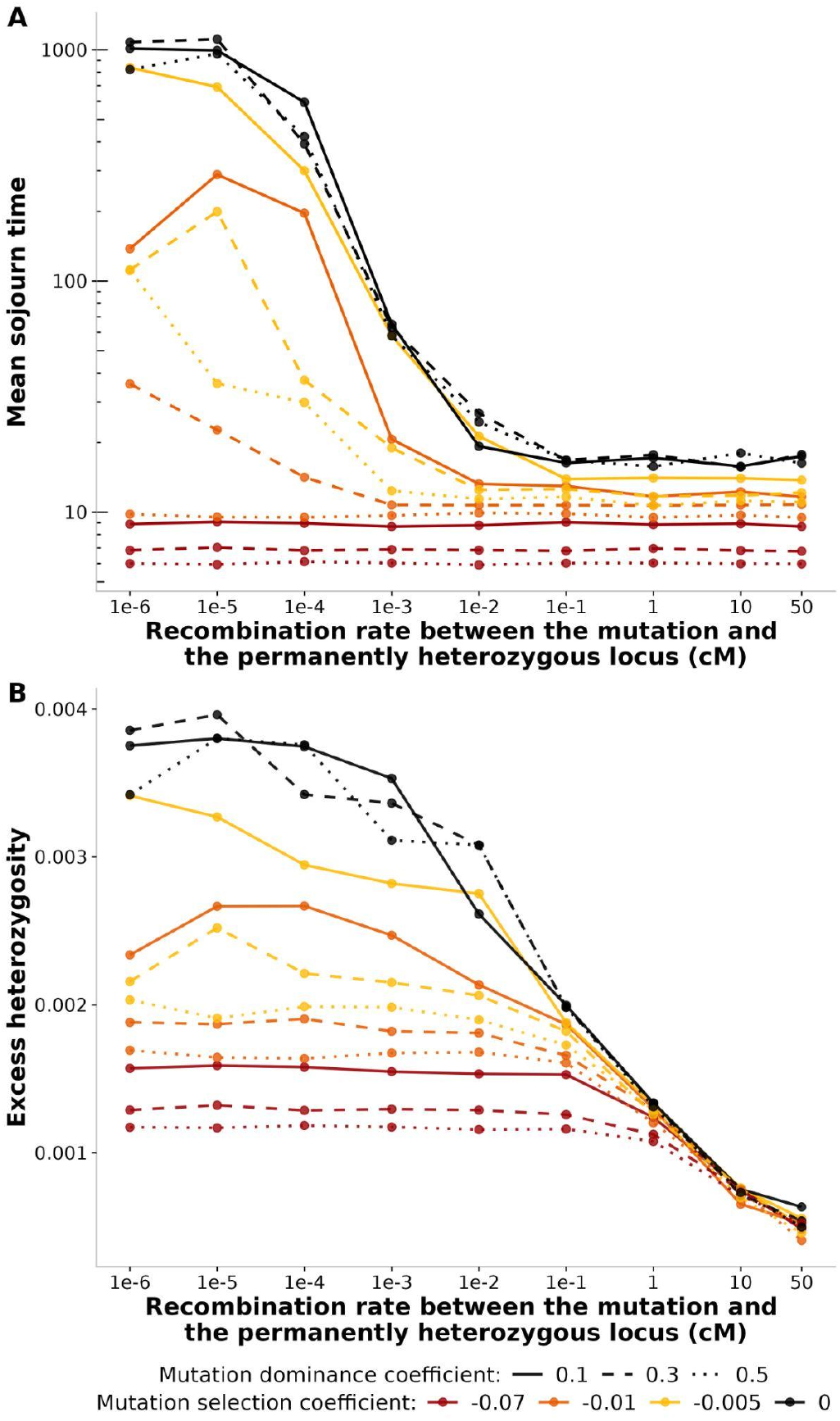
Mutation dynamics as a function of their distance from a permanently heterozygous locus. **A)** Mean mutation sojourn time as a function of their distance from a permanently heterozygous locus. Mean sojourn time across 40,000 mutations introduced individually in a population of size 1000 with a selfing rate of 0, at a distance ranging from 1 × 10⁻⁶ to 50 cM of a permanently heterozygous locus, for various selection and dominance coefficients. **B)** Mean excess heterozygosity (H_Obs_/H_Exp_ − 1) for mutations as a function of their distance from a permanently heterozygous locus, averaged over the course of the mutation lifetime and across 40,000 mutations introduced individually in a population of size 1000 with a selfing rate of 0, at a distance ranging from 1 × 10 ⁻⁶ to 50 cM of a permanently heterozygous locus, for various selection and dominance coefficients. Simulations were halted at generation 100,000 in cases where the mutation had not yet been fixed or lost.

### Permanently heterozygous loci favour pseudo-overdominance

To test whether permanently heterozygous loci facilitated the establishment of pseudo-overdominance around them, we conducted individual-based simulations under a Wright-Fisher model, tracking the evolution of a panmictic population of diploid individuals during 100,000 generations. Each individual had a single chromosome, subjected to recombination and mutation, carrying in its centre a bi-allelic, permanently heterozygous locus. For the purpose of illustration, we first focused on a single set of parameter values, with a selfing rate of σ=0, a chromosome with L=10⁷ selected sites, a population of size *N*=1000 individuals, a recombination rate *r*=1 × 10⁻⁹ events per selected site per generation corresponding to an effective density of selected sites per cM of approximately 10⁷, a mutation rate *µ*=1 × 10⁻⁸ mutations per base pair per generation, a selection coefficient of mutations *s*=-0.04 and a dominance coefficient *h*=0.3. We then also conducted sets of simulations i) with only neutral mutations (*s*=0), as a control to assess the influence of selection, and ii) without any permanently heterozygous locus, as a control to assess the influence of this type of locus on pseudo-overdominance establishment.

In simulations with a permanently heterozygous locus and deleterious mutations, the mean nucleotide diversity steadily increased along generations, going from 0 to around 2 × 10⁻⁴ in 100,000 generations (continuous orange line in Fig 2A). The mean nucleotide diversity peaked at the permanently heterozygous locus; the peak height and width increased along generations (Fig 2B), showing that partially recessive deleterious mutations accumulated around the permanently heterozygous locus. The fraction of sites with heterozygous mutations in individuals also tended towards 1 along generations (continuous orange line in Fig 2C), indicating that the partially recessive deleterious mutations were mostly heterozygous. At generation 50,000, the chromosome showed an 8 Mb block of sites in strong linkage disequilibrium with each other around the permanently heterozygous locus (Fig 2D). Linkage disequilibrium was progressively established between the sites with deleterious mutations over generations, reaching 4,000 sites in strong linkage disequilibrium in 100,000 generations (continuous orange line in Fig 2E). Hierarchical clustering indicates the presence of two complementary haplotypes composed of partially recessive deleterious mutations arranged in repulsion (Fig 2F). This indicates the existence of a pseudo-overdominant zone.

**Figure 2:**
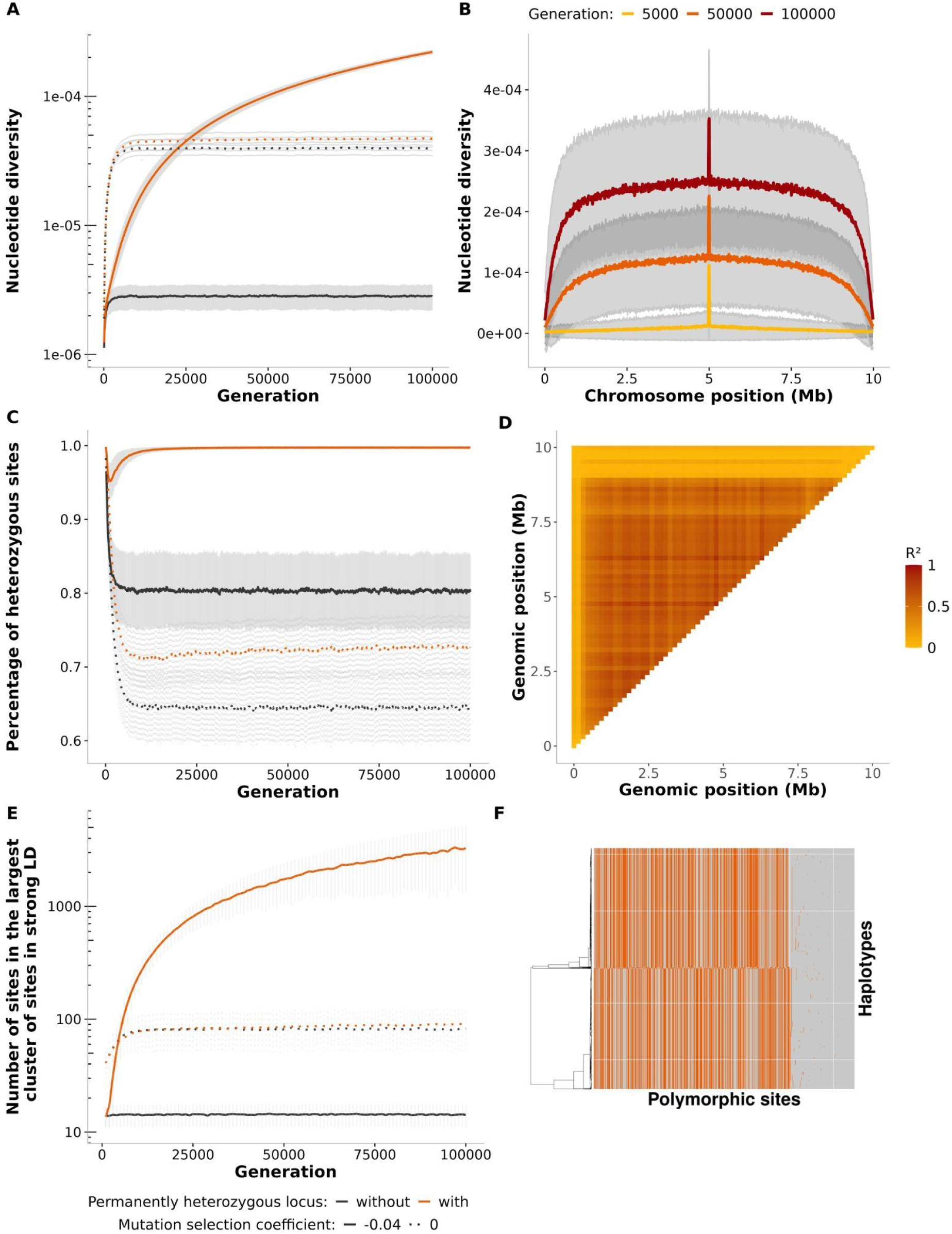
Dynamics of mutation accumulation with time and along the genome depending on their selection coefficient and of the presence of a permanently heterozygous locus. Results from the evolution of a panmictic population during 100,000 generations under a Wright-Fisher model. Each individual is diploid with a pair of chromosomes containing a central permanently heterozygous locus in some cases. Simulations used a number of selected sites (L) of 10⁷, a selfing rate (σ) of 0, a population size (*N*) of 1000, a recombination rate (*r*) of 1 × 10⁻⁹, a mutation rate (*µ*) of 1 × 10⁻⁸, and a dominance coefficient (*h*) of 0.3. **A)** Mutation accumulation over time. Mean nucleotide diversity over generations, computed every 100 generations in 100 genomes across 1000 simulations, depending on the presence of a permanently heterozygous locus (with: orange line; without: black line) and the selection coefficients of mutations (s=-0.04: continuous lines; s=0: dotted line). Standard deviation is represented in grey. The Y axis is in logarithmic scale. **B)** Partially recessive deleterious mutation accumulation around a permanently heterozygous locus. Mean nucleotide diversity computed in 100 genomes per segment of 10,000 bp in 1000 simulations with mutations with a selection coefficient of *s*=-0.04. The three lines represent the mean number of segregating mutations at three generations: 5000, 50,000, and 100,000. Standard deviation is represented in grey. **C)** Percentage of heterozygosity in individuals over time. Mean percentage of heterozygous sites per individual (among polymorphic sites) over generations, measured in 100 individuals every 100 generations in 1000 simulations, depending on the presence of a permanently heterozygous locus (with: orange line; without: black line) and the selection coefficients of mutations (s=-0.04: continuous lines; s=0: dotted line). Standard deviation is represented in grey. The Y - axis is in logarithmic scale. **D)** Linkage disequilibrium around a permanently heterozygous locus. Linkage disequilibrium map established from 100 genomes at generation 50000 with a mutation having a selection coefficient of *s*=-0.04 in the presence of a permanently heterozygous locus. **E)** Number of sites in linkage disequilibrium over generations. Number of sites in the largest cluster of sites in strong linkage disequilibrium (LD) with each other (rZ>0.95) measured from 100 individuals every 1000 generations from 1000 simulations, depending on the presence of a permanently heterozygous locus (with: orange line; without: black line) and the selection coefficients of mutations (s=-0.04: continuous lines; s=0: dotted line). Standard deviation is represented in grey. The Y -axis is in logarithmic scale. **F)** Haplotype structure across polymorphic sites when deleterious mutations having a selection coefficient of *s*=-0.04 and a permanently heterozygous locus. Each row represents an individual haplotype and each column a polymorphic site from the whole population at generation 100,000. Colours indicate allele states (grey: ancestral allele, orange: derived allele). Haplotypes are clustered based on similarity using hierarchical clustering.

In these simulations, recombination tended to generate individuals with lower fitness than average (Fig 3A) and this effect was stronger when the recombination event occurred closer to the permanently heterozygous locus (Fig 3B), as this led to offspring with a higher number of homozygous, partially recessive deleterious mutations (Fig 3C). Because the nucleotide diversity and number of sites in linkage disequilibrium increased along generations (Fig 2A,E), the mean fitness of offspring resulting from recombination decreased along generations (Figs 3A, B). The mean relative fitness of recombinant offspring declined from 1 to around 0.1 in 100,000 generations and recombination events became lethal at generation 100,000 when occurring at less than 2.5 Mb from the permanently heterozygous locus; these recombination events indeed led to homozygosity for ca. 1000 mutations (Fig 3). A pseudo-overdominant zone (i.e. arrays of recessive deleterious mutations behaving as overdominant) can thus progressively form, be maintained and extend around a permanently heterozygous locus, despite recombination, which is effectively suppressed due to the death of recombinant offspring.

In contrast, our simulations under this same set of parameter values but without any permanently heterozygous locus, and/or with only neutral mutations (*s*=0), did not show such an increase in the mean nucleotide diversity and heterozygosity nor the formation of large blocks of sites in linkage disequilibrium (black and dotted lines in Figs 2A,C,E). Instead, the nucleotide diversity and the extent of linkage disequilibrium reached each a plateau, reflecting a mutation-drift equilibrium when mutations were neutral, and a mutation-selection-drift equilibrium when they were deleterious. The absence of accumulation of polymorphic sites in linkage disequilibrium with neutral mutations confirms that drift alone cannot explain the maintenance of deleterious mutation arrays, and therefore that selection plays a role in the formation of blocks of deleterious mutations in repulsion (i.e. pseudo-overdominant zone). In addition, recessive deleterious mutations accumulate in repulsion only in the presence of a permanently heterozygous locus, confirming its role in promoting pseudo-overdominance.

### Conditions for pseudo-overdominance establishment

To further explore the impact of population genetics parameters and of the presence of a permanently heterozygous locus on the establishment of pseudo-overdominance, we simulated chromosomes containing L = 10⁵, 10⁶ or 10⁷ selected sites, with populations with sizes *N* of 1,000 or 10,000, a selfing rate σ of 0, 0.5 or 0.9, with recombination rates *r* of 1 × 10⁻⁷, 1 × 10⁻⁸, or 1 × 10⁻⁹ events per selected site per generation corresponding to an effective density of selected sites per cM of approximately 10⁵, 10⁶ or 10⁷, a mutation rate *µ* of 1 × 10⁻⁸ or 1 × 10⁻⁹ events per base pair per generation, a selection coefficient *s* of −0.01, −0.04, −0.07, −0.001, or −0.005, and a dominance coefficient *h* of 0.1, 0.3 or 0.5; we performed simulations with and without a permanently heterozygous locus and ran simulations with *s*=0 as a control. We chose these values based on estimates in natural populations (Supplementary text 2).

We considered that pseudo-overdominance was established at generation 100,000 for a given set of parameter values when the mean number of sites in the largest cluster of sites in strong linkage disequilibrium with each other (rZ>0.95), averaged over 30 replicates, was significantly higher than the corresponding mean obtained in simulations with only neutral mutations. This, together with the observation of alternance of deleterious mutations and wild-type alleles and the fitness cost of recombination (Figs 2F and 3), indeed indicate that selection promoted the formation of complementary blocks of partially recessive deleterious mutations, leading to an apparent overdominance of loci with deleterious alleles. Hereafter, we refer to such regions spanned by clusters of sites in strong linkage disequilibrium as ‘pseudo-overdominant zones’.

Overall, pseudo-overdominance emerged under a broader range of parameter combinations in the presence of a permanently heterozygous locus than in its absence. Indeed, among the 1620 conditions tested, 163 led to pseudo-overdominance with a permanently heterozygous locus, compared with only 37 without such a locus, representing a 4.4-fold increase (Figs 4A, S1 with parameters scaled by *N*, S2 with raw data). In addition, the variance in the number of sites in the pseudo-overdominant zone among replicates tended to be smaller in systems with a permanently heterozygous locus than without it, suggesting a lower influence of stochasticity in the presence of such a locus (Fig S2; 29 of the 37 conditions had a significantly lower variance with a permanently heterozygous locus than without it). Finally, the mean number of sites in linkage disequilibrium in the pseudo-overdominant zone tended to be higher in the presence of a permanently heterozygous locus than without it (Fig S1; 32 conditions out of 37 had a significantly higher mean with than without a permanently heterozygous locus). This suggests that, beyond expanding the range of conditions under which pseudo-overdominance can occur, a permanently heterozygous locus also increases the speed and extent of its establishment.

**Figure 3:**
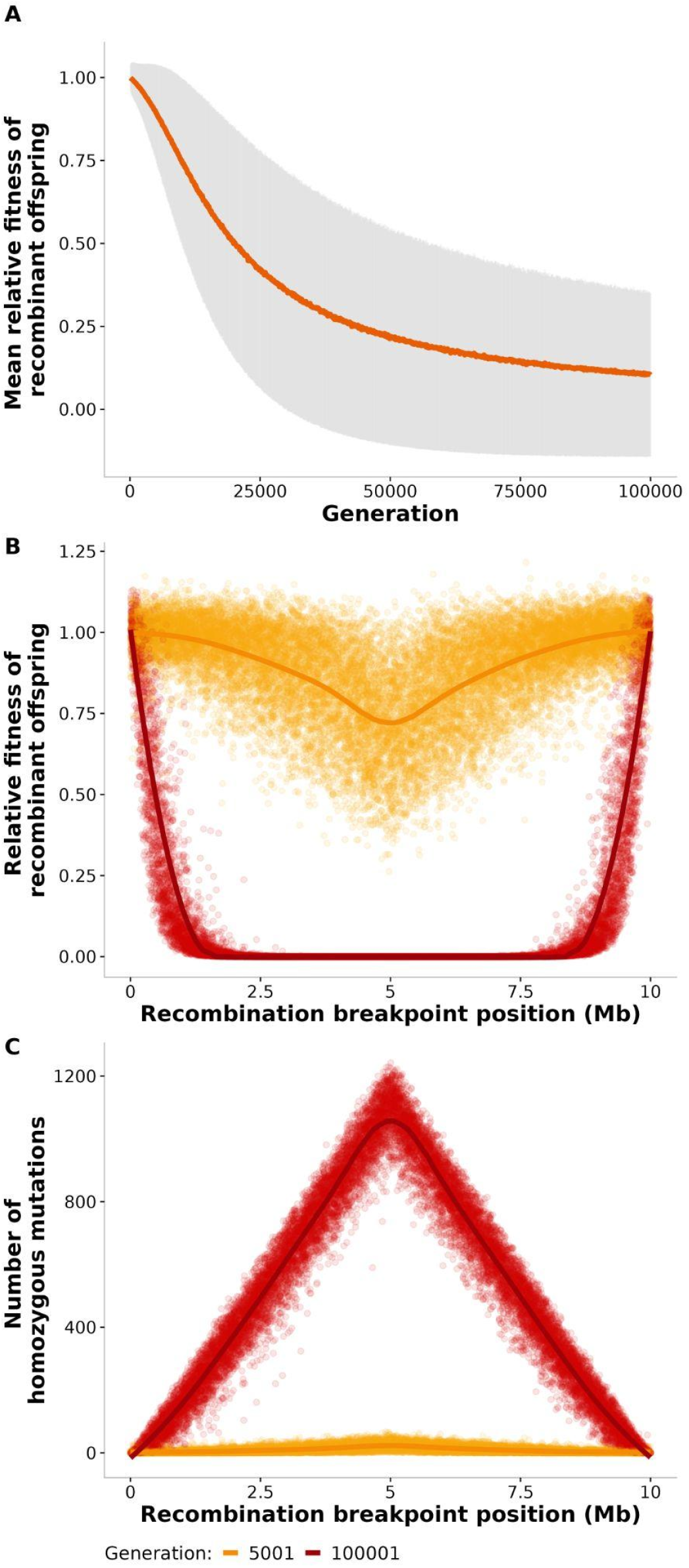
Fitness cost of recombination across the genome and over generations when pseudo-overdominance evolves. Results from 1000 simulations of the evolution of a panmictic population during 100,000 generations under a Wright-Fisher model. Each individual is diploid with a pair of chromosomes containing a central permanently heterozygous locus. Simulations used a number of selected sites (L) of 10⁷, a selfing rate (σ) of 0, a population size (*N*) of 1000, a recombination rate (*r*) of 1 × 10⁻⁹, a mutation rate (*µ*) of 1 × 10⁻⁸, a dominance coefficient (*h*) of 0.3 and a selection coefficient (*s*) of −0.04. **A)** Mean relative fitness, computed every 100 generations, of all individuals resulting from a single recombination event (i.e., resulting from one recombinant gamete generated by a single crossover in the previous generation) . **B)** Relative fitness of all individuals resulting from a single recombination event depending on the position of the recombination breakpoint, computed at generations 5,001 and 100,001. Each dot represents a recombinant individual, and the curve represents the locally estimated scatterplot smoothing. **C)** Number of homozygous sites among polymorphic sites in all individuals resulting from a single recombination event according to the position of the recombination breakpoint computed at generations 5,001 and 100,001. Each dot represents a recombinant individual, and the curve represents the locally estimated scatterplot smoothing.

**Figure 4:**
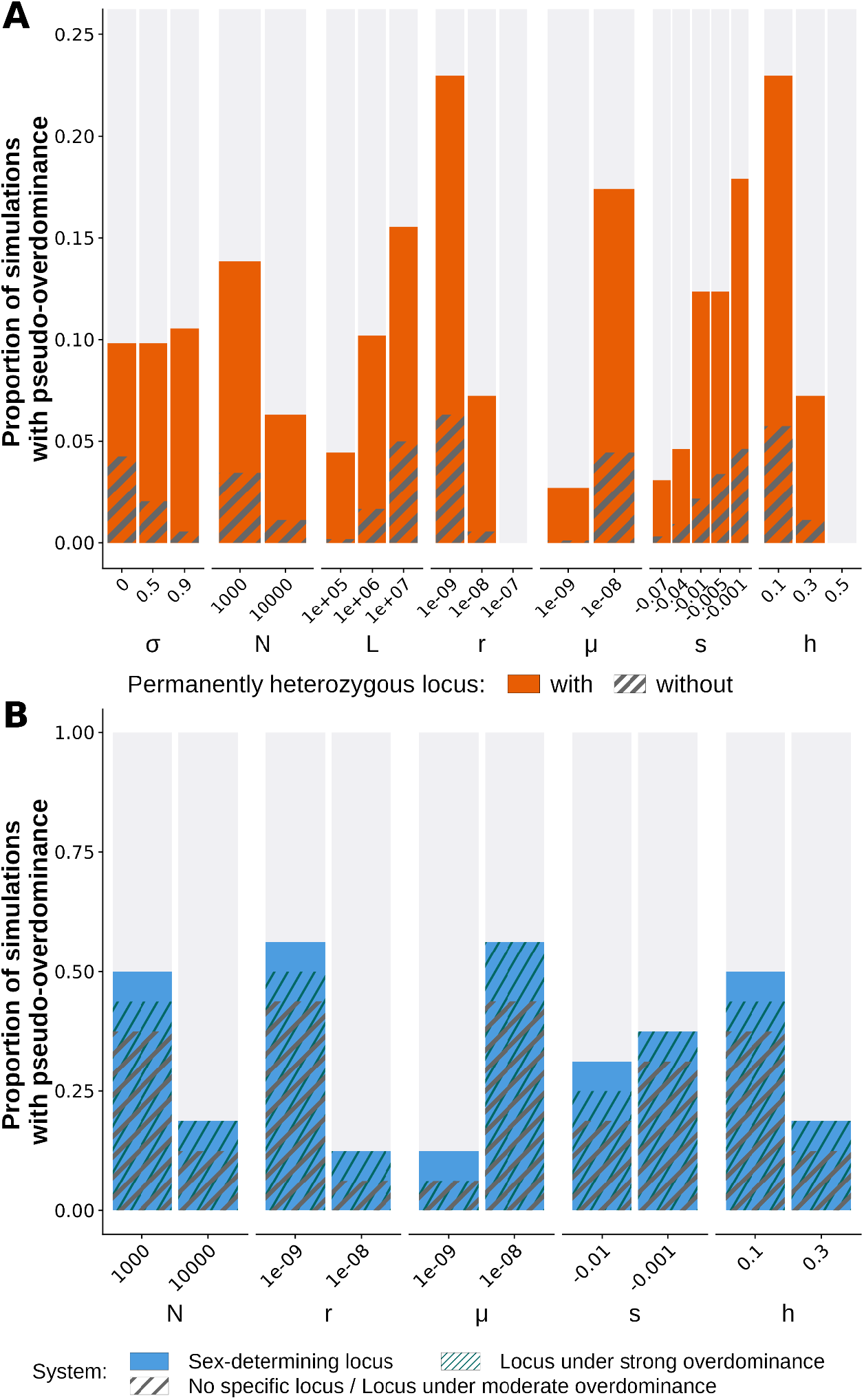
Effect of selfing rate, population size, number of sites under selection, mutation rate, recombination rate, selection and dominance coefficients of mutations and of the presence of different types of loci under balancing selection on the establishment of pseudo-overdominance. **A)** Proportion of simulations leading to pseudo-overdominance for each parameter value tested among simulations run with a selfing rate σ (0, 0.5, 0.9), a population size *N* (1000 or 10,000), a chromosome of size L (10⁵, 10⁶, 10⁷), a mutation rate *µ* (1 × 10⁻⁸, 1 × 10⁻⁹), a recombination rate *r* (1 × 10⁻⁷, 1 × 10⁻⁸, 1 × 10⁻⁹), a selection coefficient *s* (-0.01, −0.04, −0.07, −0.001, −0.005) and a dominance coefficient *h* (0.1, 0.3, 0.5), and including or not a permanently heterozygous locus in the centre of the chromosome (solid orange and dashed grey bars, respectively). **B)** Proportion of simulations leading to pseudo-overdominance for each parameter value tested among simulations run with a chromosome of size L=10⁷, a selfing rate σ=0, a population size *N* (1000 or 10,000), a mutation rate *µ* (1 × 10⁻⁸, 1 × 10⁻⁹), a recombination rate *r* (1 × 10⁻⁸, 1 × 10⁻⁹), a selection coefficient s (-0.01, −0.001) and a dominance coefficient *h* (0.1, 0.3), and including or not a sex-determining locus (solid blue) or a locus under moderate (grey dashed bar; also for no specific locus) or strong overdominance (dark green dashed bar) in the centre of the chromosome.

While selfing generally impaired pseudo-overdominance establishment, the opposite was observed around a permanently heterozygous locus, where selfing instead promoted its establishment. In the absence of a locus under balancing selection, the parameter space allowing pseudo-overdominance establishment progressively decreased with increasing selfing rates (Figs 4A and S2). Indeed, among the 1080 conditions tested, pseudo-overdominance was established in 23, 11, and 3 conditions at selfing rates of σ = 0, 0.5, and 0.9, respectively. In contrast, in the presence of a permanently heterozygous locus, pseudo-overdominance was established in a slightly larger parameter range under selfing than under outcrossing (Figs 4A and S2). Indeed, among the 1080 conditions tested with a permanently heterozygous locus, pseudo-overdominance was established in 53, 53, and 57 conditions at selfing rates of σ = 0, 0.5, and 0.9, respectively. In addition, with a permanently heterozygous locus, the mean number of sites in linkage disequilibrium within the pseudo-overdominant zone tended to increase with the selfing rate (Fig S2). In particular, 10 conditions out of 53 show a significantly higher mean number of sites in linkage disequilibrium when comparing cases with pseudo-overdominance between σ=0 and σ=0.9, 9 between σ=0.5 and σ=0.9 and 4 between σ=0 and σ=0.5, while the remaining showed no significant differences.

Overall, these results suggest that selfing generally hinders pseudo-overdominance establishment, except around a permanently heterozygous locus, where selfing can even favor it to some extent. This is likely due to several effects. First, the sojourn time and excess heterozygosity of mutations closely linked to the permanently heterozygous locus were not substantially affected by selfing (Fig S3 analogous to Fig 1). In contrast, the sojourn time and excess heterozygosity of mutations decreased, as expected, with distance to the permanently heterozygous locus with this decline becoming more pronounced as the selfing rate increased beyond 10^-1^ cM (Fig S3). Second, genetic drift is stronger under selfing, which likely explains the slightly larger condition range under which pseudo-overdominance is established with a permanently heterozygous locus under selfing, as genetic drift further facilitates the maintenance of slightly deleterious mutations.

We also found that pseudo-overdominance was more often established with a higher number of sites under selection along the chromosome (4.4 times more with L=10⁷ than L=10⁵; Figs 4A, S1 and S2). Furthermore, in agreement with previous studies (Gilbert et al. 2020; Waller 2021; Abu-Awad and Waller 2023), pseudo-overdominance was more often established with smaller population sizes (2.3 times more cases with *N*=1000 than with *N*=10000), higher mutation rates (7.7 times more cases with *µ*=1 × 10⁻⁸ than with *µ*=1 × 10⁻⁹), lower recombination rates (3.8 times more cases with *r*=1 × 10⁻⁹ than with *r*=1 × 10⁻⁸ and 0 cases with *r*=1 × 10⁻⁷), lower coefficients of dominance (3.4 times more cases with *h*=0.1 than *h*=0.3 and 0 cases with *h*=0.5) and mutations with smaller fitness effects (6.6 times more cases with *s*=-0.001 than with *s*=-0.07; Figs 4A and S1). For a particular set of parameter values, chosen as being favourable to pseudo-overdominance establishment in our simulations above (*L*=10⁷, σ=0, *N* = 1000, *µ* = 1 × 10⁻⁸, *r* = 1 × 10⁻⁹, s=-0.04, *h* = 0.3), we additionally showed that using a distribution of fitness effects for mutations instead of a single value did not substantially change our conclusions regarding pseudo-overdominance establishment, although the variance among replicates appeared higher than in the case with a fixed selection coefficient (Fig S4).

### Other overdominant loci favor pseudo-overdominance establishment

We further showed that pseudo-overdominance establishment could be facilitated around other types of loci under balancing selection with less extreme overdominance than for a permanently heterozygous locus. For this, we ran simulations with either a sex-determining locus or overdominant loci with varying degrees of heterozygote advantage, using a reduced set of parameter values to save computation time (Table 1). In the latter case, the fitness advantage of heterozygotes at the overdominant locus was set to either 0.1 (“moderate overdominance”) or 0.4 (“strong overdominance”), while the fitness gain of the two homozygotes was set to 0.0002 and 0, in both scenarios. As with a permanently heterozygous locus, we found that pseudo-overdominance emerged across a slightly broader range of parameter values around a sex-determining locus or a locus under strong balancing selection than in cases without such loci (Figs 4B, S1 and S5). Indeed, among the 32 conditions tested, 11 led to pseudo-overdominance with a sex-determining locus, 10 with a strongly overdominant locus, while only 8 led to pseudo-overdominance without such loci. This represented, under the set of parameter values tested, around 1.4 times more cases with a sex-determining locus and 1.3 more cases with a strongly overdominant locus, respectively, than without such loci. The locus under moderate overdominance did not show any detectable effect on the establishment of pseudo-overdominance under the conditions tested.

**Table 1:**
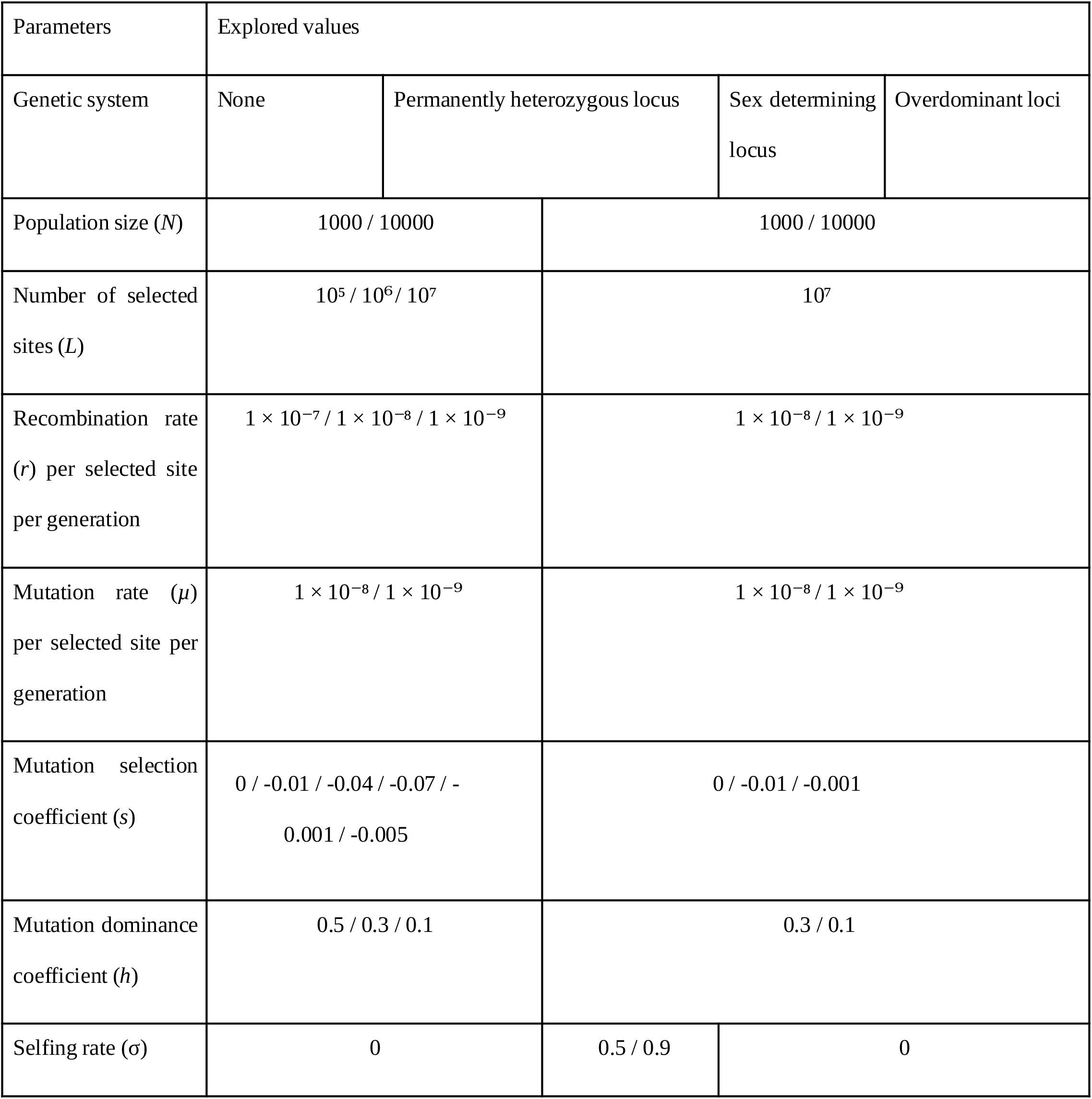
Explored parameter values in simulations.

In order to understand how pseudo-overdominance can establish in the proto-XY system (i.e., with a sex-determining locus), while XX individuals could in principle be homozygous, we performed hierarchical clustering on genotypes from simulations with the proto-XY system. We detected the presence of complementary haplotypes between proto-X and Y chromosomes, but also multiple haplotypes on the proto-X chromosome (Fig S6A and B). This means that the recessive deleterious mutations on the X chromosome were also sometimes sheltered by a complementary haplotype in XX individuals.. We additionally showed that similarly to the case with a permanently heterozygous locus (Fig 2F), two haplotypes coexisted for the case investigated without any locus under balancing selection (Fig S6C).

Overall, these results support our hypothesis that loci under balancing selection favour pseudo-overdominance, and further show that this effect intensifies with the strength of balancing selection.

### Pseudo-overdominance zones can select for true recombination suppression around permanently heterozygous loci

The strong selection against recombinant offspring shown above (Fig 3) suggests that genuine recombination suppression, such as that caused by an inversion, could be favoured in pseudo-overdominant zones, as it would avoid the production of unfit offspring via recombination. When a permanently heterozygous locus is flanked by a pseudo-overdominant region, all offspring from a non-recombining parent remain heterozygous for mutations within that region. In contrast, recombination within the pseudo-overdominant region produces some offspring carrying homozygous recessive deleterious mutations (Figs 2 and 3). This makes the offspring of non-recombining individuals fitter on average than the offspring of recombining parents, which could therefore drive the evolution of true recombination suppression in the pseudo-overdominant zone (Fig S7).

Analytical calculations showed that a recombination suppressor could have a selective advantage in a pseudo-overdominant region. We considered the simplest case, which corresponds to a recombination suppressor capturing the permanently heterozygous locus in a region of size *l* where recombination rate is *r* and every recombination event is lethal (because all recombination events lead to individual homozygous for a large number of recessive deleterious mutations; see red line in Fig 3B). In this region, recombination can be modelled as X recombination events distributed uniformly along the chromosome, with X following a Poisson distribution of parameter *rl*. Thus, the probability that there are no recombination events is P(X=0)=exp(-*rl*). Then, in the Wright-Fisher model, the advantageous allele is considered to have 1+*s* times more offspring on average than the disadvantaged allele. With this reasoning, we obtain 1+*s* = (average number of offspring with the recombination suppressor) / (average number of offspring without the recombination suppressor) = 1/exp(-*rl*) ∼ 1 + *rl* when *rl*<<1, which gives *s*≈*rl*. Here, *l* and *r* can be defined either with respect to the sites under selection only or with respect to all sites in the region, provided that their product *rl* determines the total recombination rate across the region. This provides an upper bound estimate of the selective advantage of a recombination suppressor containing a permanently heterozygous locus in a pseudo-overdominant zone; this indeed corresponds to the case in which selection against recombination is maximal, as recombination leads to lethality. With *r*= 1 × 10⁻⁹ recombination events per selected site per generation and *l*=10⁶ selected sites (i.e., a genetic distance of 0.1 cM and a density of 10⁷ selected sites / cM), the resulting selective advantage is roughly *s* = 0.001.

Simulations confirmed that a recombination suppressor could be selected for in a pseudo-overdominant zone. Due to computational constraints, this was investigated under a single set of parameter values for illustrative purposes, for which we showed above that pseudo-overdominance was established, and recombination was lethal over at least 1 Mb with 10 ⁶sites under selection i.e. spanning 0.1 cM (*L*= 10⁷, *N*=1000, σ=0, *r*=1 × 10⁻⁹, *µ*=1 × 10⁻⁸, *s*=-0.04 , *h*=0.3; Figs 2, 3, 4). We introduced a recombination suppressor in this 1 Mb encompassing the permanently heterozygous locus 1.5 × 10⁶ times at generation 100,000 and recorded the number of fixation events. As we simulated here only recombination suppressors capturing the permanently heterozygous locus, no homozygote for the non-recombining region could occur. Here, fixation was defined as the spread of the suppressor to all haplotypes carrying the permanently-heterozygous allele with which it was initially associated, corresponding to a frequency of 50% in the entire population. We observed 1,594 fixation events, significantly exceeding the 1,500 expected under neutrality (1.5 × 10 ⁶× 1/1,000; P = 0.008334, one-sided exact Poisson test). This neutral expectation assumes an initial suppressor frequency of 1/1,000 among haplotypes carrying the associated mating-type allele (i.e., half of the 2,000 chromosomes in the population of 1,000 diploid individuals). Thus, pseudo-overdominance increased suppressor fixation by 6.3% in the parameter range studied. This higher number of fixation events for a recombination suppressor in a pseudo-overdominant zone than in the neutral case (with only drift acting) confirms that the presence of a pseudo-overdominant zone facilitates the evolution of recombination suppression.

The non-recombining fragments that eventually fixed did not, on average, initially carry a lower mutational load than those that were lost (Fig S8). This indicates that recombination suppressors were selected for because they prevented the formation of unfit recombinant haplotypes and not because non-recombining fragments were simply less loaded than average (M. Nei et al. 1967; Jay et al. 2024, 2025). These results further support that pseudo-overdominance can favour the evolution of recombination suppression around permanently heterozygous loci.

## Discussion

In this study, we demonstrated that permanently heterozygous loci, sex-determining loci and loci under strong overdominance facilitate the evolution of pseudo-overdominance, which is characterized by the formation of complementary haplotypes carrying deleterious mutations in repulsion. We also found that, while selfing generally hinders the establishment of pseudo-overdominance, it can instead promote its establishment in the presence of permanently heterozygous loci. Furthermore, we showed that, once pseudo-overdominance is established around permanently heterozygous loci, selection can favour the evolution of genuine recombination suppression. The emergence of pseudo-overdominance and recombination suppression requires specific conditions (in terms of number of selected sites, recombination and mutation rates, as well as selection and dominance coefficients), and is therefore expected to occur exclusively in particular genomic contexts, such as around certain mating-type loci under selfing, where it may help explain puzzling patterns observed in these regions.

### Overdominant loci favour pseudo-overdominance establishment

Various theoretical models have investigated the conditions under which pseudo-overdominance can emerge (B. Charlesworth and D. Charlesworth 1997; Pamilo and Pálsson 1998; Zhao and B. Charlesworth 2016; Gilbert et al. 2020; Berdan et al. 2021; Waller 2021; Abu-Awad and Waller 2023; Sianta et al. 2023; Booker and Schrider 2026). These studies highlighted the impact of recombination and mutation rates on pseudo-overdominance establishment, as well as of selection and dominance coefficients, in agreement with our results. Here, we illustrated in more detail how pseudo-overdominance establishes, extends, strengthens and is maintained, especially around permanently heterozygous loci, and how it shapes haplotype structure and affects the fitness of recombinant offspring. We further demonstrated that, beyond the extreme case of permanent heterozygosity, overdominant loci more generally favour pseudo-overdominance establishment. This had been verbally suggested (Abu-Awad and Waller 2023). However, to our knowledge, pseudo-overdominance around loci under balancing selection has only been previously theoretically studied to explain the evolution of the MHC region, although not referred to as such (van Oosterhout 2009). Our results showed that the presence of a permanently heterozygous locus, a sex-determining locus or a locus under strong overdominance expands the range of conditions under which pseudo-overdominance arises. Pseudo-overdominance can indeed be established around a locus under balancing selection in systems with fewer selected sites, a higher recombination rate, a lower mutation rate, a stronger negative selection coefficient, a higher dominance coefficient and a larger population size compared to conditions enabling pseudo-overdominance without any specific loci. Pseudo-overdominant zones may involve more than two haplotypes, particularly under particular genetic systems of sex determination. In proto-XY systems, for example, pseudo-overdominance can emerge with one haplotype on the Y and multiple haplotypes on the X, such that the deleterious effects of homozygosity are not experienced by all individuals of either sex. Further investigations on the haplotypic dynamics underlying pseudo-overdominance and how these dynamics are influenced by parameter values, would therefore be valuable. We further demonstrated that the impact of an overdominant locus on the establishment of pseudo-overdominance increased with the strength of its overdominance. It remains to be investigated whether other forms of balancing selection, such as negative frequency-dependent selection, also facilitate the establishment of pseudo-overdominance, as they may also promote the maintenance of polymorphism (D. Charlesworth 2006).

We show that permanently heterozygous loci facilitate the emergence of pseudo-overdominance in their vicinity by maintaining linked mutations at intermediate frequencies. The effects of balancing selection on linked neutral loci are well documented (D. Charlesworth 2006; Kirkpatrick et al. 2010; Gao et al. 2015; Hudson 1990; Zeng et al. 2021), with loci under balancing selection altering coalescence times, and thereby increasing neutral polymorphism and linkage disequilibrium at nearby sites, in a manner analogous to population subdivision. The dynamics of partially recessive deleterious mutations near loci under balancing selection have been less extensively studied; our results support and extend previous findings (Leach et al. 1986; Glémin et al. 2001; Antonovics and Abrams 2004; Uyenoyama 2005; Llaurens et al. 2009; Lenz et al. 2016; Tezenas et al. 2023), showing that loci under balancing selection facilitate the maintenance of heterozygous recessive deleterious mutations in their flanking regions, with this effect strengthening as mutations become less deleterious and more recessive, and more tightly linked to the selected loci. Empirical studies support these theoretical predictions by providing evidence for the presence of deleterious mutations in the vicinity of permanently heterozygous loci (Stone 2004; Llaurens et al. 2009; Mena-Alí et al. 2009; Goubet et al. 2012; Le Veve et al. 2023; Guyot et al. 2025).

The mechanisms promoting the maintenance of deleterious mutations around loci under balancing selection remain, however, to be fully understood (see discussions in D. Charlesworth, 2006; van Oosterhout, 2009; Lenz *et al*., 2016; Abu-Awad and Waller, 2023). Three non-independent processes are likely involved: i) a local subdivision of the effective population size due to linkage to a locus under balancing selection may increase genetic drift and reduce the efficacy of purifying selection on deleterious mutations (Lenz et al. 2016); ii) the rarity of recombination events breaking the association between the site under balancing selection and nearby loci may prevent mutations from moving between backgrounds, which hinders the purging or fixation of mutations in an analogous way as geographical subdivision (Hudson 1990; B. Charlesworth et al. 2003; D. Charlesworth 2006; Zeng et al. 2021); iii) the linkage of sites carrying partially recessive deleterious mutations to a locus under balancing selection may favour their maintenance in the heterozygous state, a phenomenon referred to as the “sheltering effect”, also possibly extending their persistence (Masatoshi Nei 1970; van Oosterhout 2009; Lenz et al. 2016; Jay et al. 2024).

### Selfing impairs pseudo-overdominance establishment but slightly promotes it around permanently heterozygous loci

As expected, selfing generally hindered pseudo-overdominance establishment, as it favored the purge of recessive deleterious mutations, a well-known effect (Lande and Schemske 1985; Glémin 2003). However, our results show that selfing slightly promotes pseudo-overdominance establishment around a permanently heterozygous locus, likely because of two effects. First, the effect of the permanently heterozygous locus on mutations nearby still exists under selfing. Indeed, we showed that recessive deleterious mutations could be maintained close to a permanently heterozygous locus regardless of the selfing rate, consistent with theoretical results (Tezenas et al. 2023). Second, selfing increases genetic drift, thus further facilitating linkage disequilibrium building and the maintenance of slightly deleterious mutations in close linkage to the permanently heterozygous locus (Pollak 1987; Glémin 2007). These two effects combined create even more favorable conditions for pseudo-overdominance around a permanently heterozygous locus under selfing. The effect of selfing on pseudo-overdominance establishment was nevertheless weak, making it unlikely to also favor pseudo-overdominance around loci experiencing weaker overdominant selection, as higher homozygosity in their flanking regions should increase the purge of deleterious mutations. Other known effects of selfing are to promote linkage disequilibrium and haplotype differentiation at loci under balancing selection in the context of local adaptation (Nordborg 1997; Nordborg et al. 2002; Stetsenko and Roze 2022), by reducing effective recombination rates. However, this likely does not play any role in the margin of permanently heterozygous loci as these effects of selfing only act in homozygous regions, precisely because homozygosity reduces effective recombination rates.

Interestingly, all fungi with apparent recombination suppression around their mating-type loci identified so far undergo high rates of selfing, and in particular of intra-tetrad mating (De Filippo et al 2026; Jay et al 2026, Lucotte et al 2026). We could not simulate here intra-tetrad selfing (i.e. mating among the products of a given meiosis), as SLiM does not easily allow such particular crosses. However, given that intra-tetrad mating is known to maintain heterozygosity even more around a mating-type locus than classical, inter-tetrad selfing (Hood and Antonovics 2000; Tezenas et al. 2023), we can expect that pseudo-overdominance establishment should be even further facilitated under intra-tetrad mating. Indeed, intra-tetrad mating preserves heterozygosity around mating-type loci because spores of opposite mating types from the same meiosis tend to inherit different parental alleles at nearby loci, restoring heterozygosity upon fusion (Hood and Antonovics 2000). In contrast, gametes from different meioses can inherit the same parental alleles at these loci, resulting in greater loss of heterozygosity under selfing than intra-tetrad mating.

### Model relevance for natural populations

The various parameter values used here for selection and dominance coefficients, number of selected sites, their density (i.e. recombination rates between them), and for mutation rate, were chosen in agreement with previous theoretical studies (Gilbert et al. 2020; Booker and Schrider 2026), and of the order of magnitude of estimates in natural populations (Supplementary text 2; Eyre-Walker et al. 2006; Eyre-Walker and Keightley 2007; Park 2011; Manna et al. 2011; Conrad et al. 2011; Lesecque et al. 2012; Agrawal and Whitlock 2012; Duchen et al. 2013; Henn et al. 2015; Stapley et al. 2017; Halldorsson et al. 2019; Rifkin et al. 2021; Y. Wang and Obbard 2023; Bergeron et al. 2023; Di and Lohmueller 2024; Kyriazis and Lohmueller 2024; Mrnjavac et al. 2025). The parameter ranges span the transition between conditions favourable and unfavourable to the establishment of pseudo-overdominance, thereby capturing the most relevant region of the parameter space. Estimates of key parameters from natural populations, identified as drivers of pseudo-overdominance evolution (Supplementary text 2), suggest that conditions favouring pseudo-overdominance can be met in some species and specific genomic regions, although its establishment ultimately depends on the combination of parameter values. However, a key challenge in assessing whether natural parameter values allow pseudo-overdominance is that, once these conditions are met, pseudo-overdominance actually emerges and alters the apparent recombination rates and estimates of selection and dominance coefficients per site.

Other conditions not tested here could likely facilitate pseudo-overdominance in natural populations. A higher number of alleles at the locus under balancing selection and a higher ploidy are, for instance, expected to facilitate deleterious mutation maintenance and thus favour pseudo-overdominance establishment (D. Charlesworth 2006; van Oosterhout 2009; Booker and Schrider 2026). Similarly, demographic processes that increase genetic drift, such as population structure and bottlenecks, may facilitate the establishment of pseudo-overdominance, potentially broadening the conditions under which it arises in nature.

Some simplifying assumptions of our model, most of them used in previous models on pseudo-overdominance (e.g. Gilbert et al. 2020; Abu-Awad and Waller 2023; Booker and Schrider 2026), differ from conditions likely to be encountered in natural populations and may affect the conditions under which pseudo-overdominance evolves. The heterozygous advantage values tested for the overdominant loci were chosen to be high compared to known estimates (Satta et al. 1994; Yasukochi and Satta 2013), to avoid the loss of one or the other allele in the presence of deleterious mutations in the condition tested. In addition, our model assumes that sites under selection are evenly spaced along the chromosome, which may facilitate the evolution of pseudo-overdominance. Indeed, the presence of large neutral regions between sites under selection could disrupt the build-up of linkage disequilibrium among these sites, thereby hindering the emergence and extension of pseudo-overdominant regions. We also model uniform recombination and mutation rates and all mutations having the same selective and dominance coefficient. Nevertheless, simulating a distribution of fitness coefficients also led to pseudo-overdominance establishment in the favorable conditions tested. Simulating more realistic scenarios is challenging, as this would require a substantially more complex model and estimation of key parameters from natural populations remain limited. Overall our goal was not to reproduce the whole complexity and the exact genomic architecture of a specific organism, but instead to identify key parameters that influence the establishment of pseudo-overdominance.

### Empirical evidence of pseudo-overdominance around loci under balancing selection

In genomes, most regions with low recombination rates do not exhibit the elevated diversity expected under pseudo-overdominance; instead, they typically show reduced diversity, most likely due to Hill-Robertson interference (Begun and Aquadro 1992; Cutter and Payseur 2013; B. Charlesworth and Campos 2014; Badouin et al. 2017; Becher et al. 2020). Nevertheless, in line with our predictions regarding the footprint and emergence of pseudo-overdominance, studies in fish, *Drosophila*, butterflies, zebra finches, plants and humans have identified genomic regions characterized by apparent overdominance, high genetic diversity and low recombination rates, that may in fact be pseudo-overdominant zones (Frydenberg 1963; van Oosterhout 2009; Hedrick et al. 2016; Schou et al. 2017; Knief et al. 2017; Becher et al. 2020; Gilbert et al. 2020; Leitwein et al. 2021; Jay et al. 2021; Salson et al. 2025). In some cases (Frydenberg 1963; Becher et al. 2020), it has been suggested that a locus under balancing selection might be present in these regions, which may have shaped their evolution. To our knowledge, the MHC is the only locus under balancing selection for which the establishment of pseudo-overdominance, though not explicitly referred to as such, has been proposed to explain the surrounding patterns of high polymorphism organized into haplotype blocks (van Oosterhout 2009).

In addition, many well-characterized loci under balancing selection are also surrounded by regions structured into haplotype blocks, often associated with a load of partially recessive deleterious mutations, and may thus be affected by pseudo-overdominance. Classic examples include plant and vertebrate disease resistance genes, hemoglobin genes in mammals, sex-determining regions and permanently heterozygous loci (Judelson et al. 1995; Stone 2004; Llaurens et al. 2009; Mena-Alí et al. 2009; Lenz et al. 2016; Llaurens et al. 2017; B. Wang et al. 2019; Hartmann, Duhamel, et al. 2021; Berdan et al. 2022; Le Veve et al. 2023, 2024; Guyot et al. 2025; Moya et al. 2025). These may just be the result of suppressed recombination and associated Hill-Robertson effects but they could also result from pseudo-overdominance. For instance, in the fungi *Schizothecium tetrasporum* and *Podospora anserina*, the mating-type locus is surrounded by a non-recombining region spanning approximately 1 Mb, but the proximal or evolutionary causes of this recombination suppression remain unresolved. These regions are characterized by high heterozygosity, the presence of mutations disrupting coding sequences, an experimentally measured sheltered load, infrequently detected recombination events, an absence of chromosomal rearrangement and a clear haplotype structure (Contamine et al. 1996; Grognet and Silar 2015; Idnurm et al. 2015; Hartmann, Duhamel, et al. 2021; Vittorelli et al. 2023; Grognet et al. 2025; Guyot et al. 2025; De Filippo et al. 2026). This is consistent with the existence of pseudo-overdominance around mating-type loci as pseudo-overdominance leads to selection against recombinant haplotypes, generating signatures of recombination suppression while recombination events still occur. It would therefore be interesting to determine whether recombination is genuinely suppressed in such genomic regions where no recombinants have been detected despite the lack of chromosomal rearrangements (Branco et al. 2018; Guyot et al. 2025; De Filippo et al. 2026). In addition, recombination suppression around mating-type loci has been so far only found in fungal organisms with an exclusively diploid phase (Jay et al. 2024; Booker and Schrider 2026; De Filippo et al. 2026), which is a prerequisite for pseudo-overdominance to evolve. Balanced lethal systems have also been reported around mating-type loci, in fungi and oomycetes, with different lethal alleles linked to the alternative mating-type alleles, in particular in anther-smut *Microbotryum* fungi (Branco et al. 2018; Thomas et al. 2003; Hood and Antonovics 2000; Judelson et al. 1995). In sex chromosomes, some studies have reported that not only the Y chromosome does not recombine, but also the X chromosome, often following an expansion of recombination suppression around peri-centromeric regions, with initially low recombination rates (Rifkin et al. 2021; Moraga et al. 2025; Carlson et al. 2017), which reminds of the pseudo-overdominance mechanism described in the present study. This altogether suggests that pseudo-overdominance may be more prevalent than previously appreciated. Progress in sequencing should allow us to detect genomic regions displaying signatures of pseudo-overdominance (Gilbert et al. 2020). Our work may help pinpoint pseudo-overdominant zones in genomes (see supplementary text 3).

### Pseudo-overdominance can lead to selection for recombination suppression

Previous studies have shown that pseudo-overdominance could emerge in regions with low recombination rates (Sturtevant and Mather 1938; Gilbert et al. 2020; Berdan et al. 2021; Abu-Awad and Waller 2023; B. Charlesworth 2024) and that strong pseudo-overdominance prevented the evolution of increased recombination rates (Palsson 2002). Here, we show that, in the presence of a permanently heterozygous locus, pseudo-overdominance can promote the selection for true recombination suppression. Indeed, recombination between complementary haplotypes in pseudo-overdominant zones generates unfit haplotypes. Therefore, recombination suppressors, such as chromosomal inversions, could be favoured in those regions because they prevent the formation of unfit, recombinant haplotypes. The idea has been verbally suggested for sex and mating-type chromosomes, but without any mention of pseudo-overdominance (Branco et al. 2017). For illustration purposes, we focus here on a rather extreme but plausible case corresponding to a recombination suppressor capturing 10⁶selected sites in a pseudo-overdominant zone with a density of 10⁷ selective sites / cM, mimicking an inversion size of about 10 Mb if we consider that 10% of sites are under selection (see estimates from natural population in Supplementary text 2). Ten mega-base inversion falls within the range of inversion sizes documented across organisms; for example, inversions ranging from 1.5 to 43.8 Mb have been reported in deer mice (Harringmeyer and Hoekstra 2022), and inversions exceeding 10 Mb have also been documented in humans (Porubsky et al. 2022). Under these conditions, our analysis provides an upper bound estimate of the selective advantage of s=0.001 for a recombination suppressor in a pseudo-overdominant zone containing a permanently heterozygous locus. For reference, the majority of beneficial mutations in genomes have been estimated to have similarly small selection coefficients (Eyre-Walker and Keightley 2007), with for example 98% of the beneficial mutations in humans having a *s* < 0.0005 (Huber et al. 2017).

More generally, the selective advantage of a recombination suppressor in a pseudo-overdominant zone depends on how many offspring it prevents from becoming homozygous for partially recessive deleterious mutations and on the cost of being homozygous for the mutations contained within the non-recombining fragment, i.e., the recombination rate, the size of the fragment, the number of mutations, and their selection and dominance coefficients. However, because selection for recombination suppression is stronger when recombination is higher, whereas pseudo-overdominance emerges more readily when recombination is lower, the parameter space in which pseudo-overdominance both evolves and promotes recombination suppression is likely narrow.

Nevertheless, even in the absence of any locus under balancing selection, or with a locus under a balancing selection less strong than for a permanently heterozygous locus, selection for recombination suppression may also be expected in pseudo-overdominant zones, because recombination also produces unfit haplotypes in these cases. Indeed, non-recombining parents produce a higher proportion (50%) of the fittest individuals (fully heterozygous), whereas recombining parents, due to the production of recombinant haplotypes, generate fewer fully heterozygous offspring.

Roze (2021) showed that selection is expected to favor higher recombination rates as the rate of partially recessive deleterious mutations increases. In contrast, our results indicate that, under certain conditions, an increased deleterious mutation rate can promote the emergence of pseudo-overdominance, which may in turn select for reduced recombination. This apparent discrepancy likely arises because the model developed by Roze (2021) did not focus on complex selective interference among multiple linked deleterious mutations that can give rise to pseudo-overdominance, a phenomenon that emerges only under relatively restrictive conditions. Mechanisms that limit mating between gametes carrying the same haplotype, such as disassortative mating, may also be favoured when pseudo-overdominance to limit the production of offspring homozygous for partially recessive deleterious mutations (Jay et al. 2021).

## Conclusion

Our results prompt new avenues for understanding pseudo-overdominance, recombination suppression and sheltered load around loci under balancing selection. Importantly, the fact that pseudo-overdominance arises only within a restricted range of parameter values indicates that this phenomenon is likely to evolve only occasionally, such as in regions under balancing selection, and with low recombination rates, and possibly under selfing. This pattern aligns with empirical observations, where genetic signatures compatible with pseudo-overdominance are detected in genomes, but only in specific genomic regions and particular organisms. This has possibly important implications for understanding the evolution of mating-type chromosomes, but also for multi-allelic incompatibility systems, regions under balancing selection genome-wide and perhaps even sex chromosomes. Future research focusing on empirical validations of our model could provide further insights into its evolutionary implications.

## Methods

### Model

We simulated a population of *N* diploid individuals, randomly mating with a selfing rate σ, each carrying a pair of chromosomes with L selected sites, under a Wright-Fisher model with a genomic recombination rate *r*, a mutation rate *µ*, and mutation selection and dominance coefficients *s* and *h*, respectively. Fitness was calculated multiplicatively, with each mutation conferring a fitness effect of 1 + *s* when homozygous and 1 + *hs* when heterozygous. Relative fitness determined the probability of an individual being selected for reproduction. Note that parents were sampled randomly with replacement, such that self-fertilization was permitted even with σ=0. SLiM version 4.2.2 was used for all simulations (Haller and Messer 2023). The parameter values explored are given in Table 1.

In some simulations (Table 1), we added at the centre of the chromosome a permanently heterozygous bi-allelic locus controlling mating compatibility at the haploid stage (i.e. mimicking a mating-type locus), a sex-determining locus controlling mating compatibility at the diploid stage (similar to SRY in a proto-XY system), or a locus under overdominant balancing selection (with a fitness gain for heterozygous at this locus being set to be 0.1 or 0.4 while the fitness gain for each of the homozygous were respectively set to 0.0002 and 0 in both cases).

### Deleterious mutation dynamics around a permanently heterozygous locus

To evaluate how permanent heterozygosity at a locus influences the fate of nearby mutations, we analyzed the trajectories of mutations introduced individually at varying genetic distances from a permanently heterozygous locus in a population of 1000 individuals. Distances ranged from 1 × 10 ⁻⁶ to 50 cM (specifically: 1 × 10⁻⁶, 1 × 10⁻⁵, 1 × 10⁻⁴, 1 × 10⁻3, 1 × 10⁻Z, 0.1, 1, 10, and 50 cM).

Dominance coefficients (*h*) were set to 0.1, 0.3, or 0.5, selection coefficients (*s*) varied from 0 to −0.07 (specifically: 0, −0.005, −0.01, −0.07), and selfing rate (σ) were set to 0, 0.5 or 0.9. For each combination of parameters, 40,000 mutations were simulated. For each mutation, we recorded: its sojourn time and the mean excess heterozygosity corresponding to the average of (H _Obs_/H_Exp_ − 1) over the course of its lifetime, where H_Obs_ refer to the observed heterozygosity and H_Exp_ to the heterozygosity from Hardy-Weinberg expectations (i.e. 2*f*(1–*f*), where *f* is the mutation frequency). Simulations were terminated at generation 100,000 if the mutation had neither fixed nor been lost by that point.

### Illustration of pseudo-overdominance

For a subset of parameters values (*L*=10⁷, *N*=1000, σ=0, *h*=0.3, *s* =-0.04 or *s* =0, *r*=1×10⁻⁹, and *μ*=1×10⁻⁸), and under conditions with or without a permanently heterozygous locus, we recorded the nucleotide diversity of 100 genomes and the mean percentage of heterozygous sites per individual (among polymorphic sites) averaged across 100 individuals. These metrics were computed every 100 generations over 100,000 generations, across 1000 simulation replicates. The nucleotide diversity was also calculated from a sample of 100 genomes in 10 kb sliding windows along the chromosome at generations 5000, 50000 and 100000 in 1000 simulation replicates. VCFtools (Danecek et al. 2011) was used to calculate the linkage disequilibrium coefficient (rZ) between all loci along the genome in 100 individuals, sampled every 1000 generations over the course of 100,000 generations, across 1000 replicate simulations. The number of sites in the largest cluster of sites in strong linkage disequilibrium with each other of sites (rZ>0.95) was counted for each of the 1000 simulations every 1000 generations. A linkage disequilibrium map was generated from a single simulation at generation 100,000 by averaging rZ values across pairs of 150,000 bp genomic regions along the chromosome. A full classical output from SLiM was also produced at generation 100,000 and clustering was performed using hierarchical clustering based on Jaccard distances calculated from the presence/absence of polymorphic sites, with the method used in (Berdan et al. 2021). As a control, the same simulations and analyses were run for s=0. In addition, the fitness of all individuals carrying a single chromosome that had undergone a single recombination event in the previous generation was recorded every 100 generations over 100,000 generations, in 1000 simulation replicates. The number of homozygous mutations, the recombination breakpoint position and the fitness of the individuals resulting from a single recombination event were also recorded at generations 5001 and 100,001 in 1000 simulation replicates.

### Sensitivity analysis

For all combinations of parameter values (Table 1), 30 replicate simulations were run during 100,000 generations. We additionally ran simulations with mutation selection coefficients drawn from a gamma distribution with shape parameter 0.2 and mean –0.04 (setting other parameters to L=10⁷, N = 1000, σ=0, µ = 1 × 10⁻⁸, r = 1 × 10⁻⁹, h = 0.3, and including or not a permanently heterozygous locus). For each simulation, the mean number of sites in the largest cluster of sites in strong linkage disequilibrium with each other (rZ>0.95) was counted at generation 100,000 using VCFtools (Danecek et al. 2011) on a sample of 200 genomes from 100 individuals. T-tests with the Bonferroni correction for multiple testing were used to compare, for each set of parameter values (Figs S2 and S5), the mean number of sites in the largest cluster of sites in strong linkage disequilibrium with each other at generation 100,000 to the corresponding one in simulations with only neutral mutations (for *s*=0, 90 replicates were used). We used t-tests with the Bonferroni correction for multiple testing to compare the mean number of sites in the largest cluster of sites in strong linkage disequilibrium with each other at generation 100,000, between the conditions i) with and without a permanently heterozygous locus or ii) with and without selfing in cases with a permanently heterozygous locus; when both sets of conditions allowed pseudo-overdominance. F-tests were performed to compare the variance of the number of sites in the largest cluster of sites in strong linkage disequilibrium with each other between conditions with and without a permanently heterozygous locus when both sets of conditions allowed pseudo-overdominance with the Bonferroni correction for multiple testing. Additionally, for two replicate simulations with a sex-determining locus and one with no specific locus, hierarchical clustering was performed at generation 100,000, based on Jaccard distances calculated from the presence/absence of polymorphic sites, with the method used in (Berdan et al. 2021), for the following set of parameter values: L=10⁷, *N*=1000, σ=0, *h*=0.3, *r*=1 × 10⁻⁹, *µ*=1 × 10⁻⁸ and *s*=-0.01.

### Recombination suppressor introduction

In each of the 30 simulations previously run with a permanently heterozygous locus, L=10⁷, *N*=1000, σ=0, *h*=0.3, *r*=1 × 10⁻⁹, *µ*=1 × 10⁻⁸ and *s*=-0.04 a recombination suppressor of 1 Mb (capturing 10⁶ sites under selection) was then introduced 50,000 times at generation 100,001 centred around the permanently heterozygous allele of one randomly chosen genome. The number of recombination suppressor reaching fixation was then counted from each of the 1,500,000 simulations (within the subpopulation of chromosomes bearing the permanently heterozygous allele contained in the non-recombining fragment initially introduced). This large number of simulations enables us to reliably estimate the fixation probability of the recombination suppressor, which is expected to be on the same order of magnitude as that of a neutral mutation in a population size of 500 i.e 1/1000 ≈ 0.001. To approximate a fixation probability of 0.001 within ±5 % at 95 % confidence, at least 1.5 × 10⁶ simulations are needed. A one-sided exact Poisson test was used to test whether the observed number of fixation events of the recombination suppressor was significantly higher than expected under neutrality; fixation is meant here within the subpopulation of chromosomes bearing the permanently heterozygous allele contained in the non-recombining fragment initially introduced, i.e. in half the total population size. In all simulations, we also counted the initial number of mutations in each randomly chosen non-recombining fragment. A Wilcoxon rank-sum test was used to assess whether the number of mutations differed between fixed and non-fixed non-recombining fragments.

### Plot and statistical analyses

Plot and statistical analyses were performed using R software v 4.3.2 (R Core Team 2020).

### Codes and data availability

The code used to generate data and figures are available at https://gitlab.com/Louguyot/article_guyot_2025. Data will be uploaded on Zenodo upon manuscript publication.

## Acknowledgements

We thank Amandine Veber and Diala Abu-Awad for discussions, as well as Janis Antonovics and referees for valuable feedback on the manuscript. LG acknowledges a MICROBES mobility grant from the Paris-Saclay University. This work has been supported as part of the France 2030 program "ANR-11-IDEX-0003” to LG, by the European Research Council (ERC) grants EvolSexChrom (832352) and DEGENSEX (101262718) to TG and a Human Frontier Science Program (HFSP) fellowship to PJ.

## Author contribution

Conceptualization: TG, PJ and LG. Formal analysis: LG. Investigation: LG and PJ. Supervision: TG and PJ. Visualization: LG. Writing: LG, PJ and TG. Writing – review & editing: LG, TG, PJ.

## Supplementary Figures, Tables and text

**Supplementary Figure 1:**
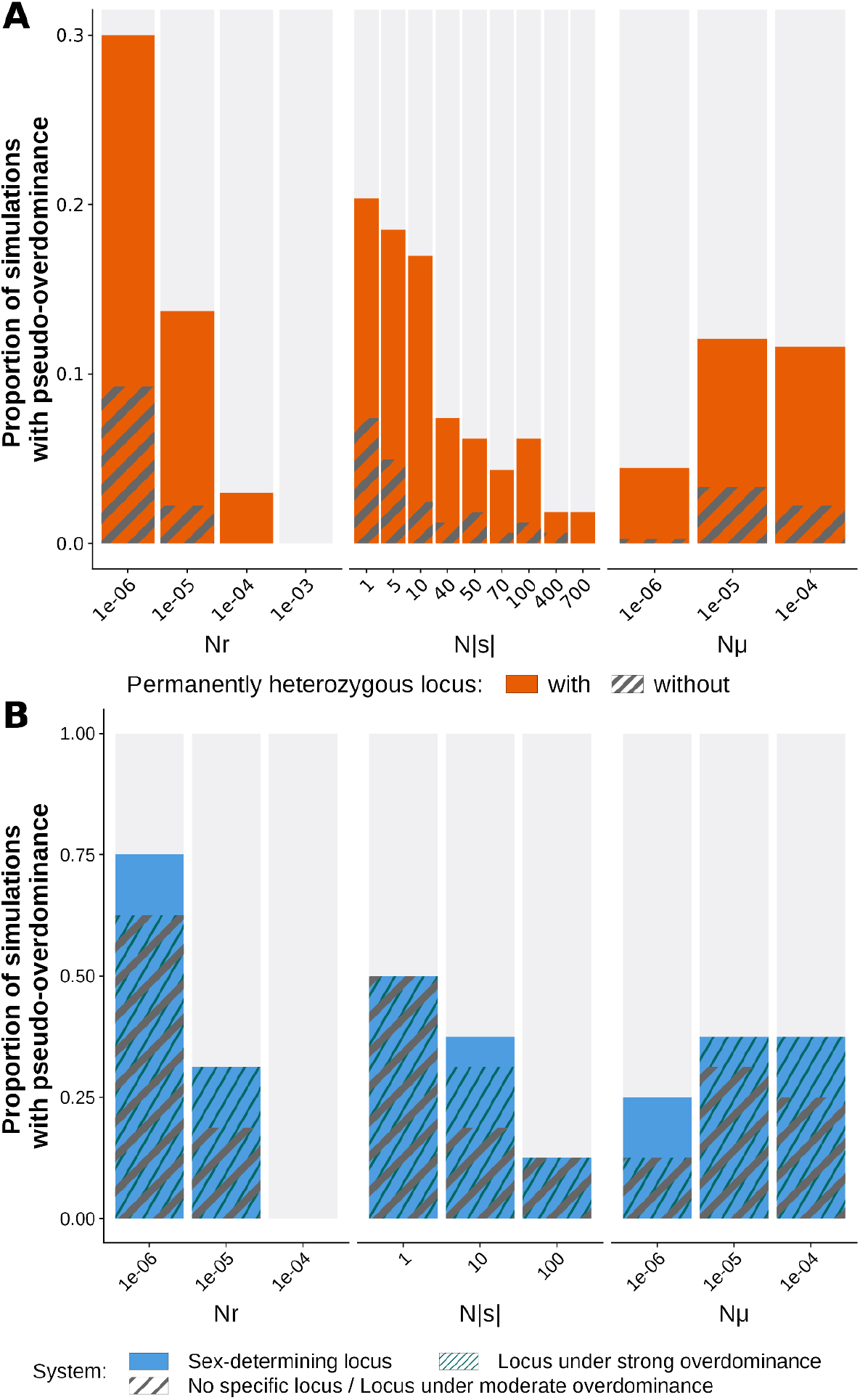
Effect of the product of the population size with the recombination rate, selection coefficients of mutations and mutation rate and of the presence of different types of loci under balancing selection on the establishment of pseudo-overdominance. A) Proportion of simulations leading to pseudo-overdominance for each product represented among simulations run with a selfing rate σ (0, 0.5, 0.9), a population size *N* (1000 or 10,000), a chromosome of size L (10⁵, 10⁶, 10⁷), a mutation rate *µ* (1 × 10⁻⁸, 1 × 10⁻⁹), a recombination rate *r* (1 × 10⁻⁷, 1 × 10⁻⁸, 1 × 10⁻⁹), a selection coefficient *s* (-0.01, −0.04, −0.07, −0.001, −0.005) and a dominance coefficient *h* (0.1, 0.3, 0.5), and including or not a permanently heterozygous locus in the centre of the chromosome (solid orange and dashed grey bars, respectively). B) Proportion of simulations leading to pseudo-overdominance for each product represented among simulations run with a chromosome of size L=10⁷, a selfing rate σ=0, a population size *N* (1000 or 10,000), a mutation rate *µ* (1 × 10⁻⁸, 1 × 10⁻⁹), a recombination rate *r* (1 × 10⁻⁸, 1 × 10⁻⁹), a selection coefficient s (-0.01, -0.001) and a dominance coefficient *h* (0.1, 0.3), and including or not a sex-determining locus (solid blue) or a locus under moderate (grey dashed bar; also for no specific locus) or strong overdominance (dark green dashed bar) in the center of the chromosome.

**Supplementary Figure 2:**
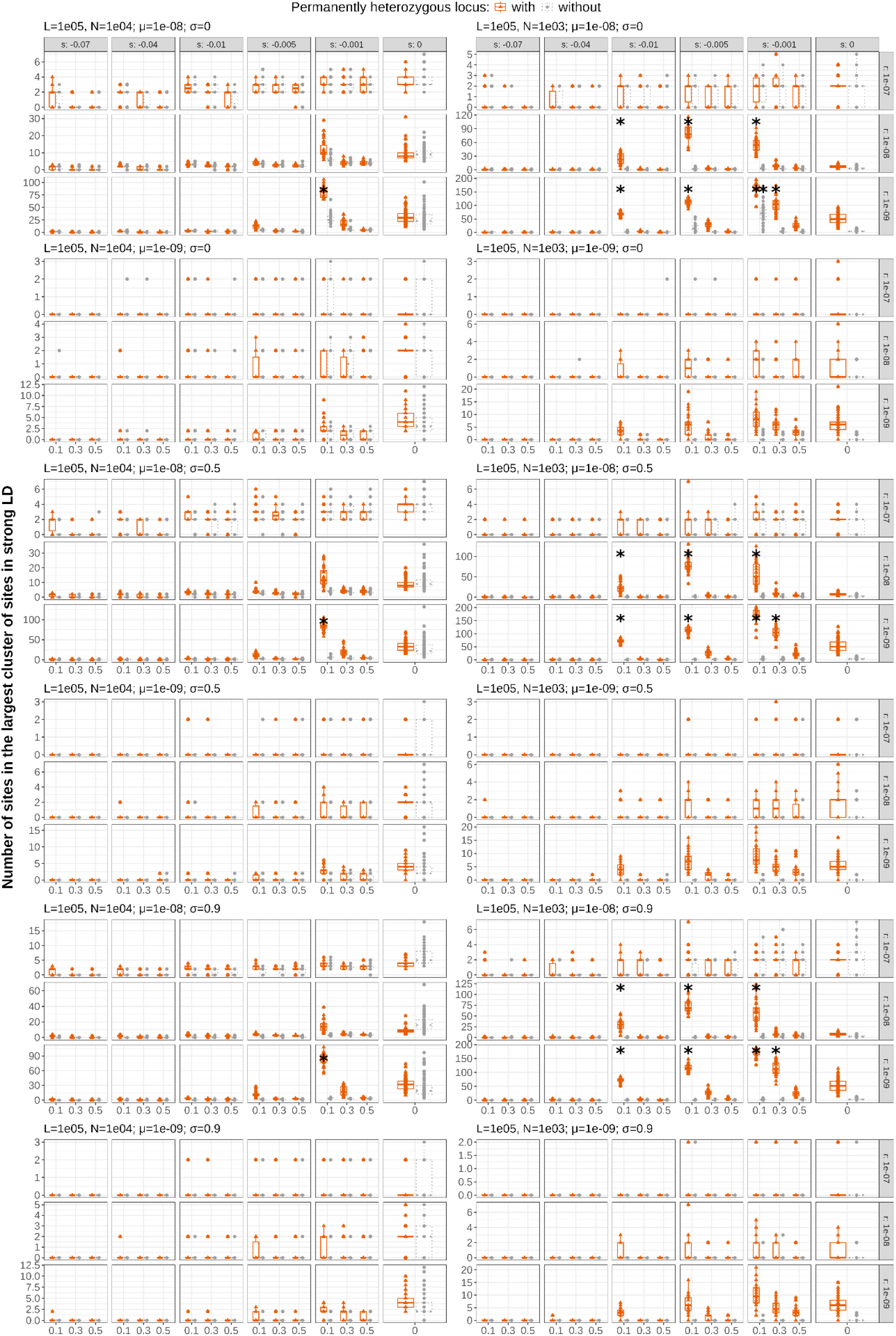
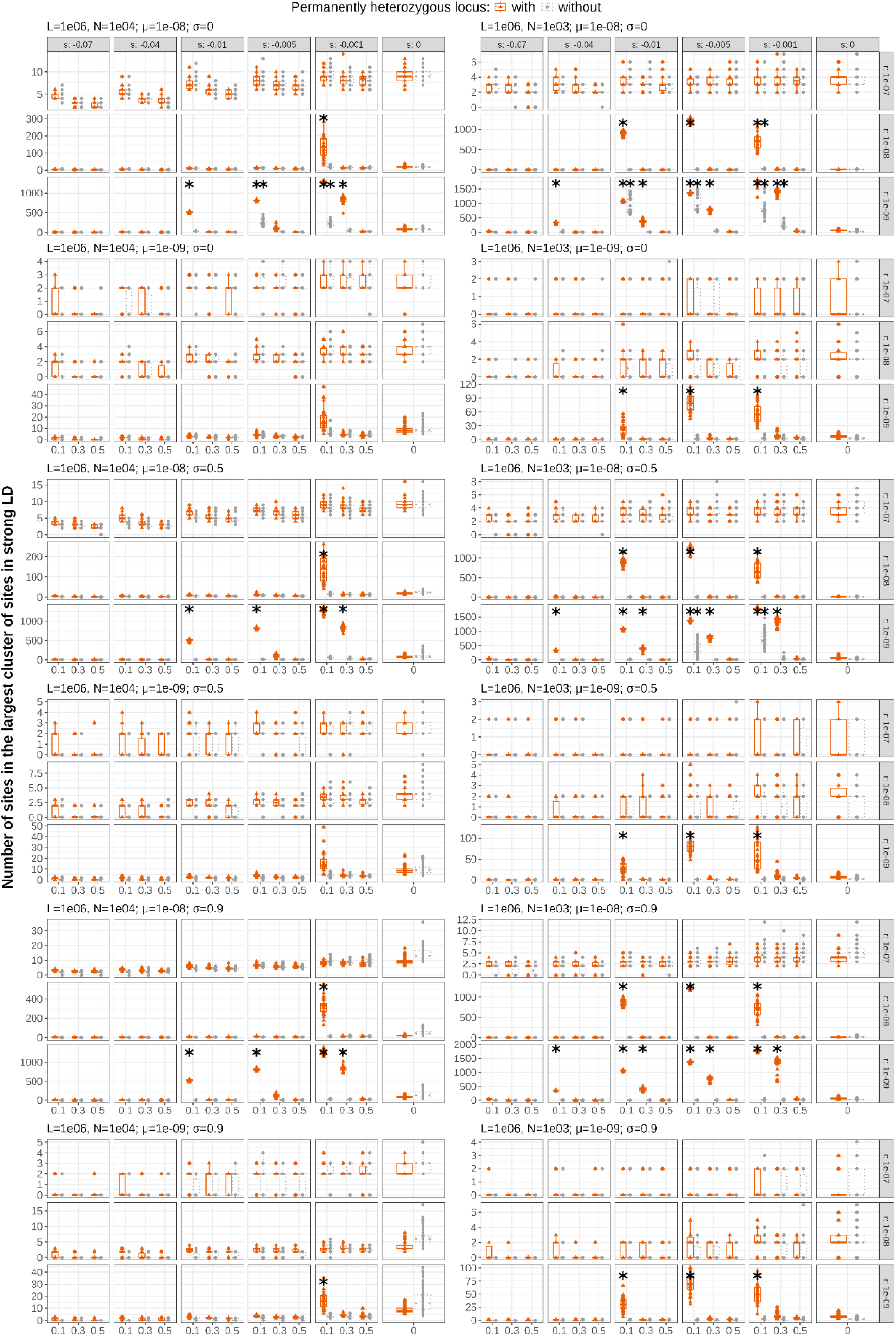
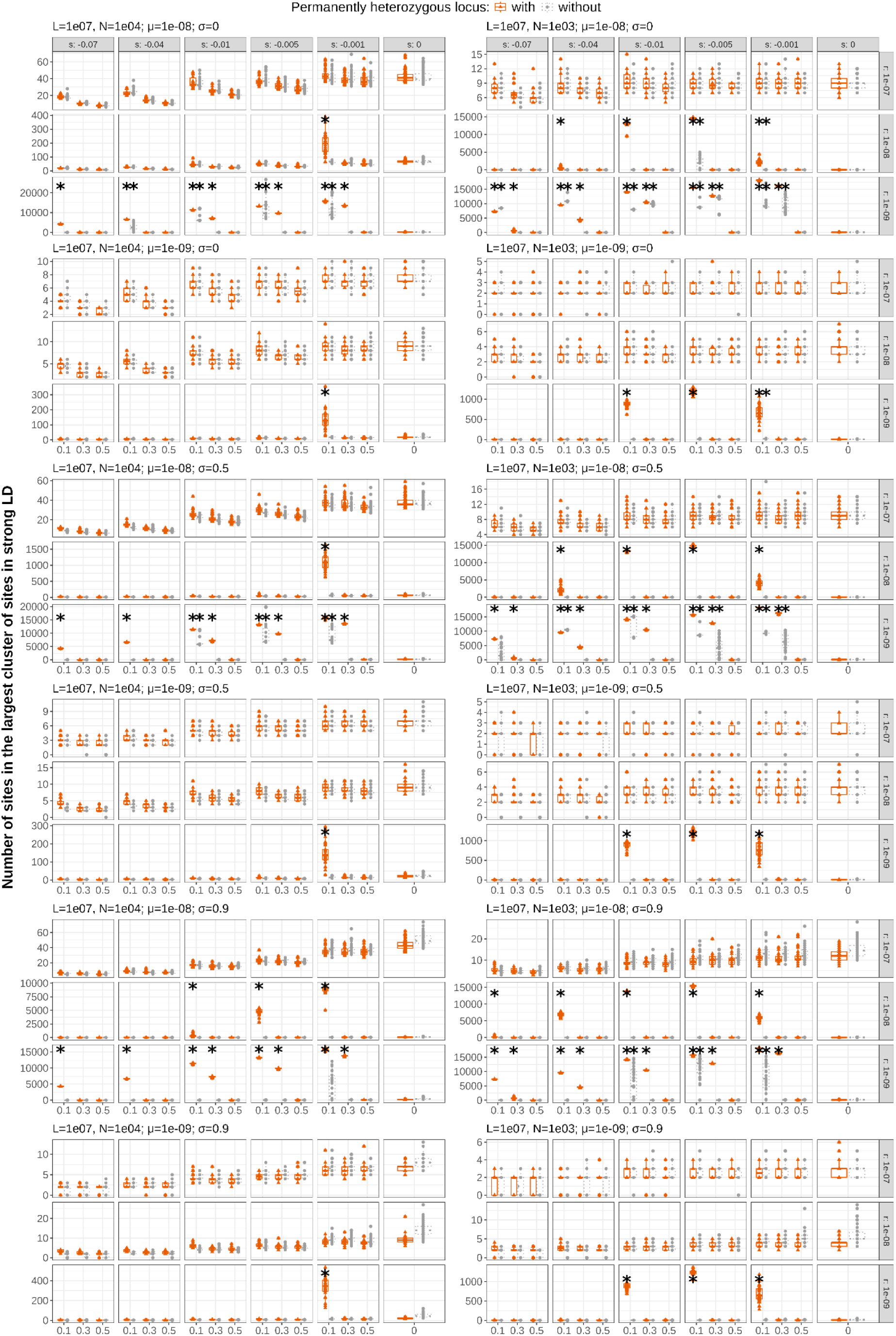
Effect of number of sites, population size, selfing rate, mutation rate, recombination rate, selection and dominance coefficients of mutations and of the presence of a permanently heterozygous locus on the establishment of pseudo-overdominance. Number of sites in the largest cluster of sites in strong linkage disequilibrium (LD) with each other (rZ>0.95) at generation 100,000 in 30 replicates of simulations (expect for *s*=0, for which 90 replicates were performed), run with various combinations of parameters: number of selected sites *L* (10⁷, 10⁶, 10⁵), selfing rate σ (0, 0.5, 0.9), population size *N* (1000 or 10,000), mutation rate *µ* (1 × 10⁻⁸, 1 × 10⁻⁹), recombination rate *r* (1 × 10⁻⁷, 1 × 10⁻⁸, 1 × 10⁻⁹), selection coefficient *s* (-0.01, −0.04, −0.07, −0.001, -0.005, 0) and dominance coefficient *h* (0.1, 0.3, 0.5). The orange boxplot on the left and the grey boxplot on the right correspond to simulations with and without a permanently heterozygous locus, respectively. The corresponding number of selected sites, population size, mutation rate and selfing rate are indicated at the top of each graph. Stars correspond to parameter values for which the mean number of sites in the largest cluster of sites in strong LD across replicates was significantly higher than the mean for the same set of parameters but with *s*=0 (t-tests with p-value corrected with Bonferroni method for multiple tests). Boxplots show the median (central line), the 25th and 75th percentiles (box limits), and whiskers extending to 1.5× the interquartile range.

**Supplementary Figure 3:**
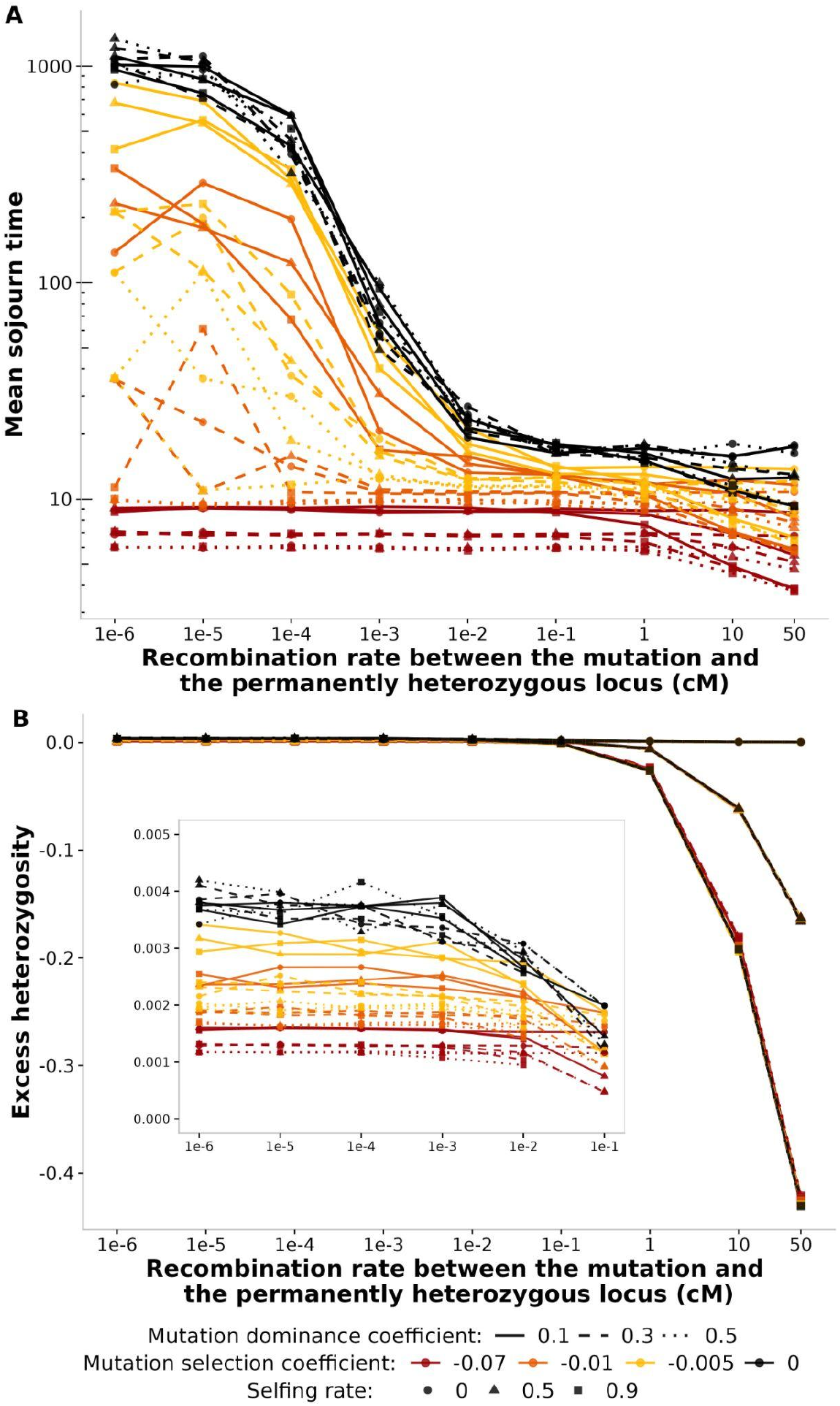
Mutation dynamics as a function of the selfing rate and their distance from a permanently heterozygous locus. A) Mean mutation sojourn time as a function of their distance from a permanently heterozygous locus. Mean sojourn time across 40,000 mutations introduced individually in a population of size 1000, at a distance ranging from 1 × 10⁻⁶ to 50 cM of a permanently heterozygous locus, for various selfing rate and selection and dominance coefficients. B) Mean excess heterozygosity (H_Obs_/H_Exp_ − 1) for mutations as a function of their distance from a permanently heterozygous locus, averaged over the course of the mutation lifetime and across 40,000 mutations introduced individually in a population of size 1000 with a selfing rate of 0, at a distance ranging from 1 × 10 ⁻⁶ to 50 cM of a permanently heterozygous locus, for various selection and dominance coefficients. Simulations were halted at generation 100,000 in cases where the mutation had not yet been fixed or lost.

**Supplementary Figure 4:**
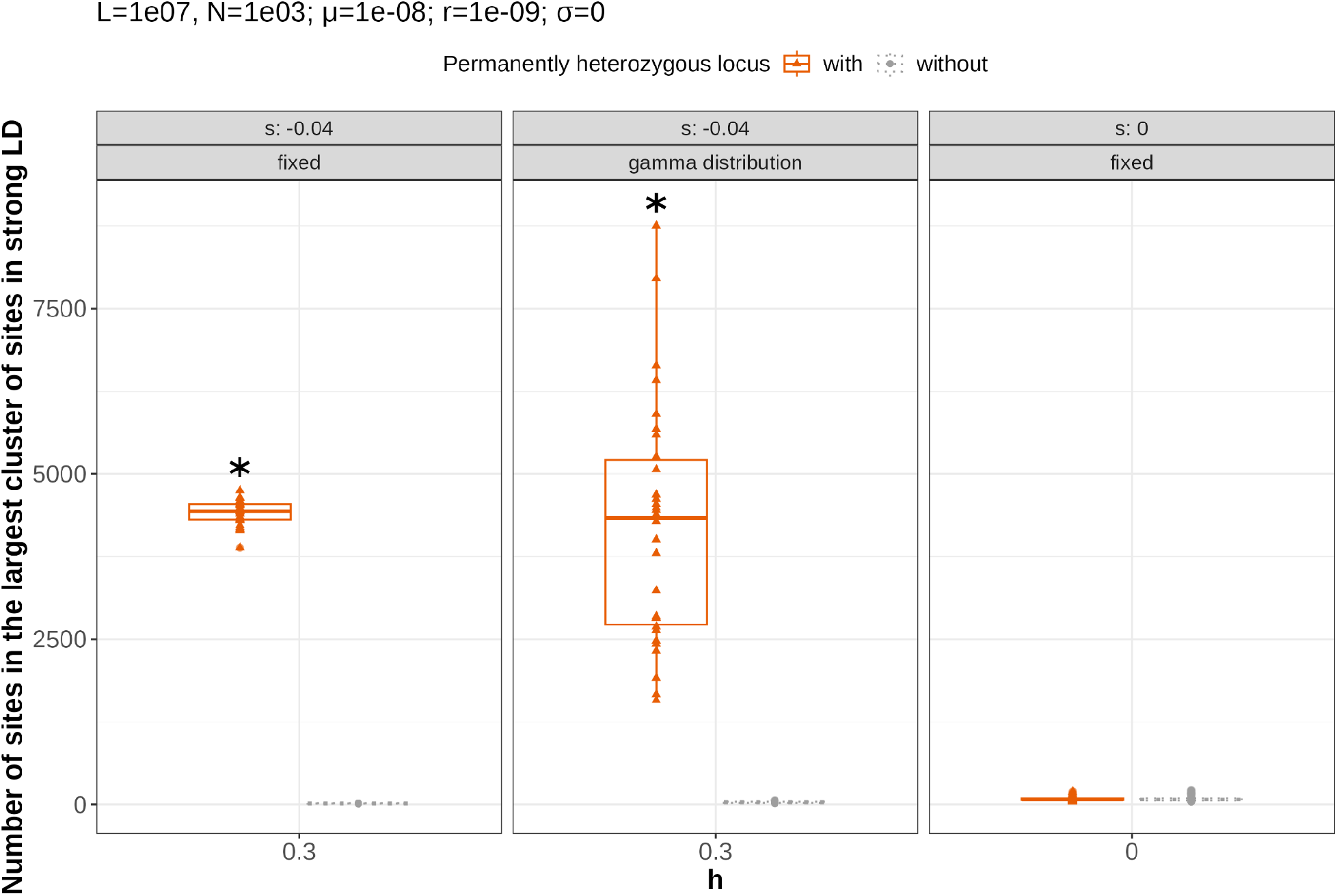
Effect of a fitness distribution on the establishment of pseudo-overdominance. Number of sites in the largest cluster of sites in strong linkage disequilibrium (LD) with each other (rZ>0.95) at generation 100,000 in 30 replicates of simulations (except for *s*=0, for which 90 replicates were performed). Simulations were run with either a fixed selection coefficient of −0.04 or a gamma distribution with shape parameter 0.2 and mean −0.04. Other parameters were set to: number of selected sites *L*=10⁷, selfing rate σ=0, population size *N* = 1000, mutation rate *µ* = 1 × 10⁻⁸, recombination rate *r* = 1 × 10⁻⁹, and dominance coefficient *h* = 0.3. The orange boxplot on the left and the grey boxplot on the right correspond to simulations with and without a permanently heterozygous locus, respectively. Stars correspond to parameter values for which the mean number of sites in the largest cluster of sites in strong LD across replicates was significantly higher than the mean for the same set of parameters but *s*=0 (t-tests with p-value corrected with Bonferroni method for multiple tests). Boxplots show the median (central line), the 25th and 75th percentiles (box limits), and whiskers extending to 1.5× the interquartile range.

**Supplementary Figure 5:**
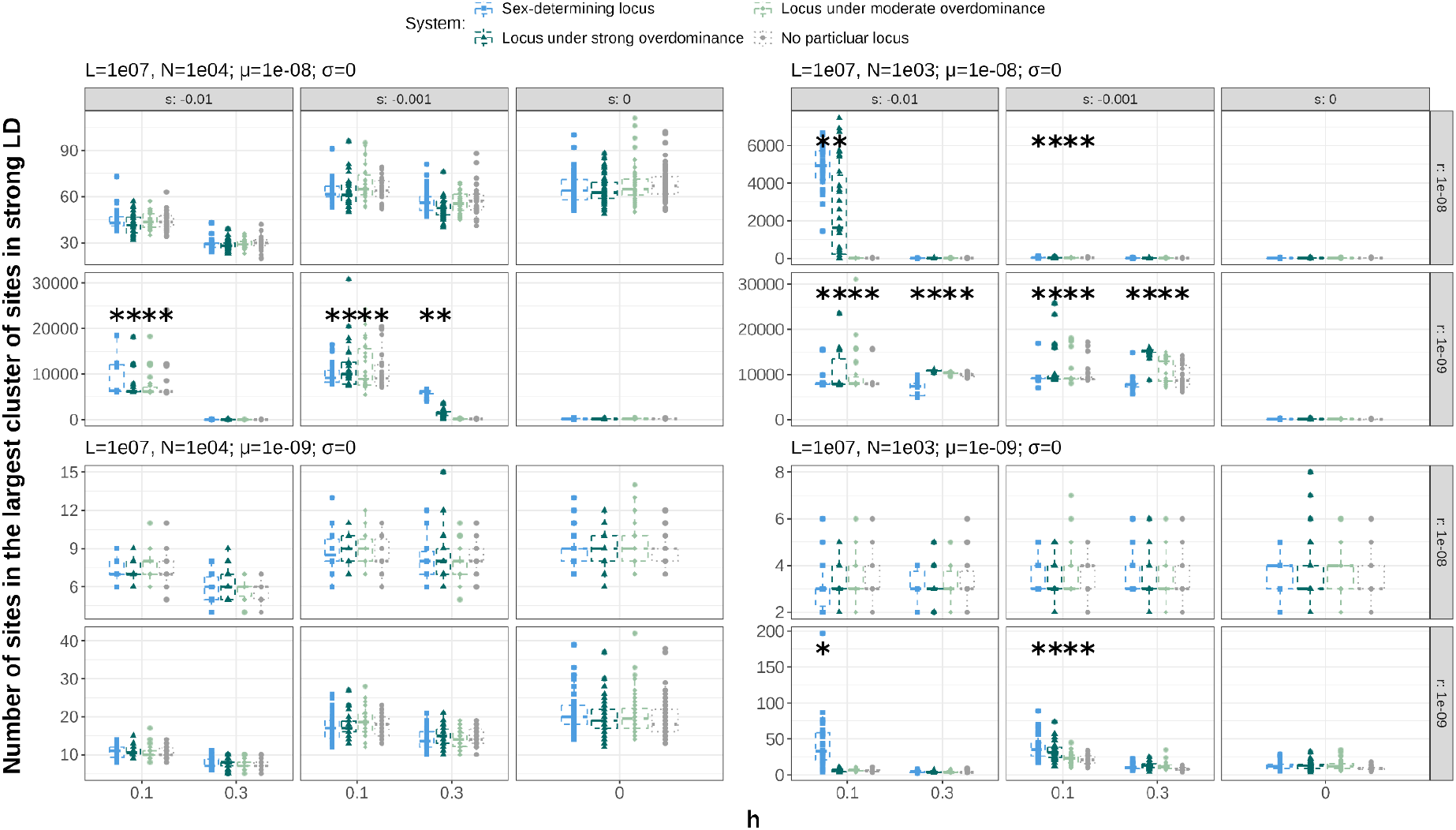
Effect of population size, mutation rate, recombination rate, selection and dominance coefficient of mutations and genetic system (sex-determining locus, overdominant loci or none) on the establishment of pseudo-overdominance. Number of sites in the largest cluster of sites in strong linkage disequilibrium (LD) with each other (rZ>0.95) at generation 100,000 in 30 replicates of simulations (expect for *s*=0, for which 90 replicates were performed) run with various combinations of parameters: population size *N* (1000 or 10,000), mutation rate *µ* (1 × 10⁻⁸, 1 × 10⁻⁹), recombination rate *r* (1 × 10⁻⁸, 1 × 10⁻⁹), selection coefficient *s* (-0.01, −0.001, 0) and dominance coefficient *h* (0.1, 0.3) in a proto-XY system or including a locus under moderate or strong overdominance or not. The corresponding number of selected sites (set to 10⁷), population size, mutation rate and selfing rate σ (set to 0) are indicated at the top of each graph and for each set of parameter values, the boxplots from left to right correspond respectively to simulations in a system including a sex-determining locus (blue), a locus under strong overdominance (dark green, with a fitness gain for heterozygotes at this locus being set to be 0.4 while the fitness gains for the two homozygotes were set to 0.0002 and 0), a locus under moderate overdominance (light green, with a fitness gain for heterozygotes at this locus being set to be 0.1 while the fitness gains for the two homozygotes were set to 0.0002 and 0), or no particular locus (grey). Stars correspond to parameter values for which the mean number of sites in the largest cluster of sites in strong LD across replicates was significantly higher than the mean for the same set of parameters (i.e. *N*, *µ*, *r,* genetic system) but *s*=0 (t-tests with p-value corrected with Bonferroni method for multiple tests). Boxplots show the median (central line), the 25th and 75th percentiles (box limits), and whiskers extending to 1.5× the interquartile range.

**Supplementary Figure 6:**
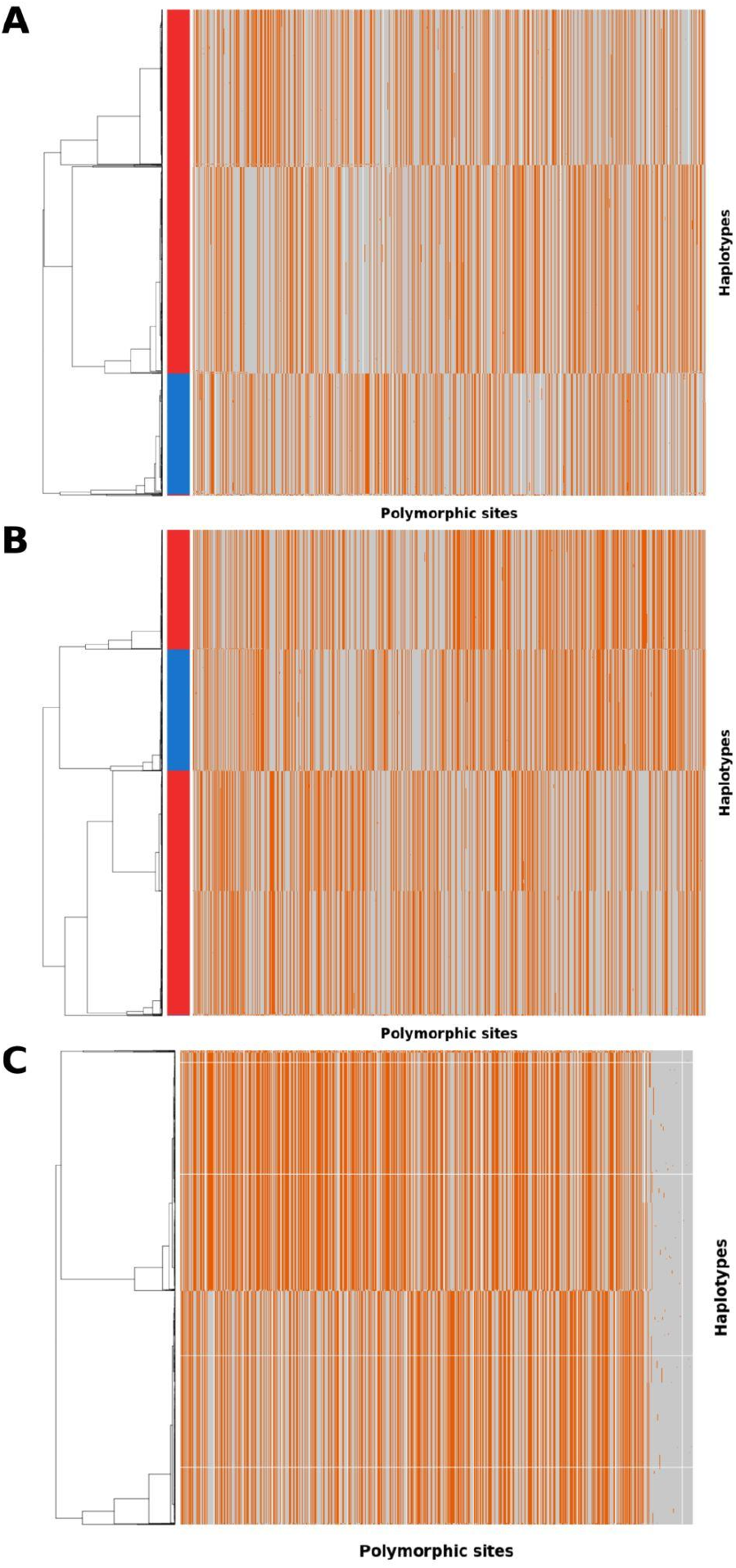
Haplotype structure when pseudo-overdominance around a sex-determining locus or no specific locus. Haplotype structure across polymorphic sites at generation 100,000 of a simulation run under a Wright-fisher model with a number of selected site L=10⁷, a selfing rate σ=0, a population size *N*=1000, a mutation rate *µ*=1 × 10⁻⁸, a recombination rate *r*=1 × 10⁻⁹, mutations with a selection coefficient *s*=-0.01 and dominance coefficient *h*=0.3. Two replicates A-B) are represented for simulations run with a sex determining locus (proto-X and Y chromosomes), one with C) no particular locus. For all plots each row represents an individual haplotype and each column a polymorphic site from the whole population at generation 100,000. Colored bars on the left indicate X chromosomes in red and Y chromosomes in blue (for A-B), color on the plots indicate allele states (grey: ancestral allele, orange: derived allele). Haplotypes are clustered based on similarity using hierarchical clustering.

**Supplementary Figure 7:**
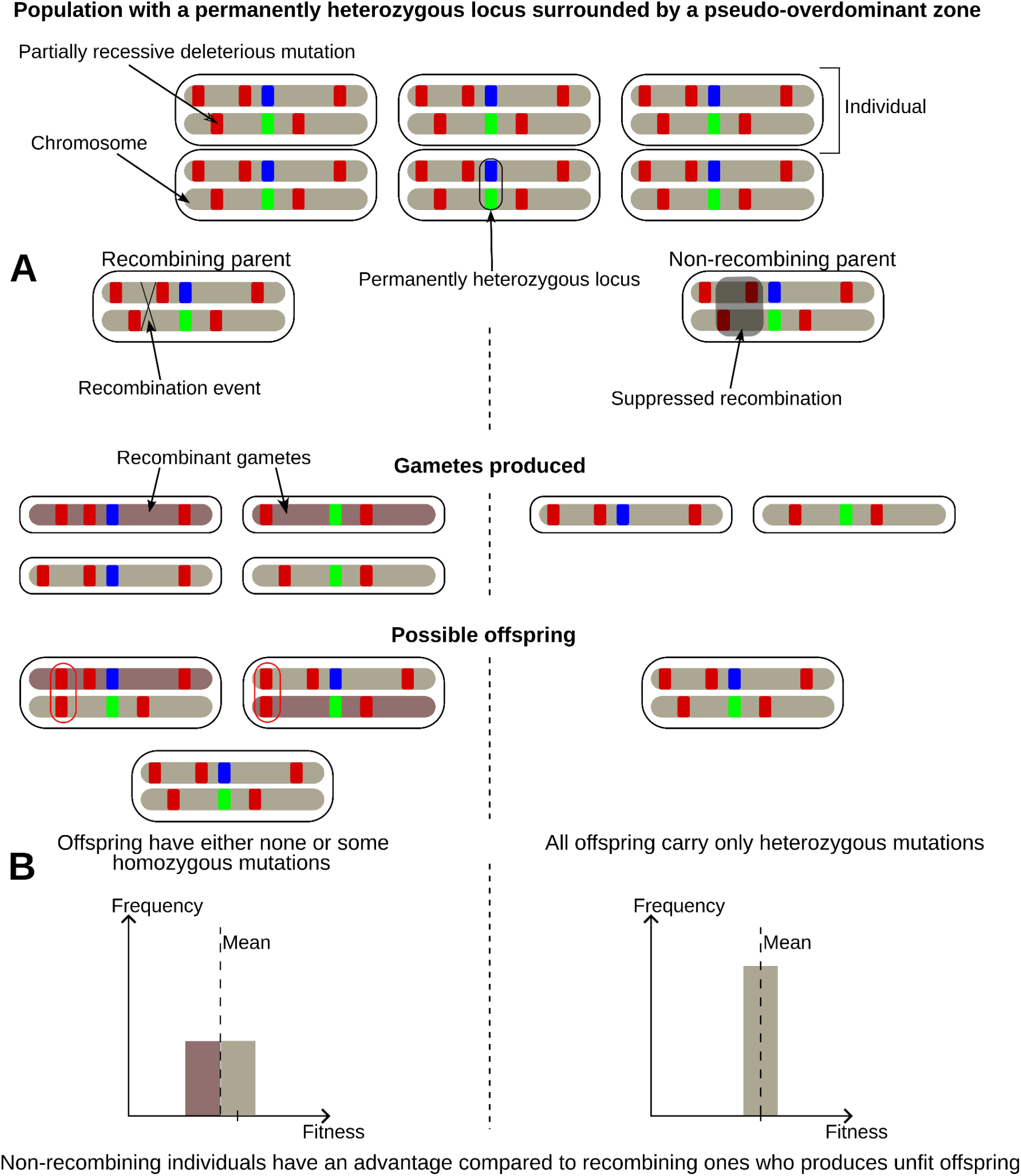
Illustration of the selective advantage of recombination suppression in a pseudo-overdominant zone surrounding a permanently heterozygous locus. A) Illustration of the production of offspring with homozygous recessive deleterious mutations by recombinant parents (at left) but not by non-recombinant parents (at right). B) Distribution of fitness in progenies of recombinant parents (at left) and non-recombinant parents (at right).

**Supplementary Figure 8:**
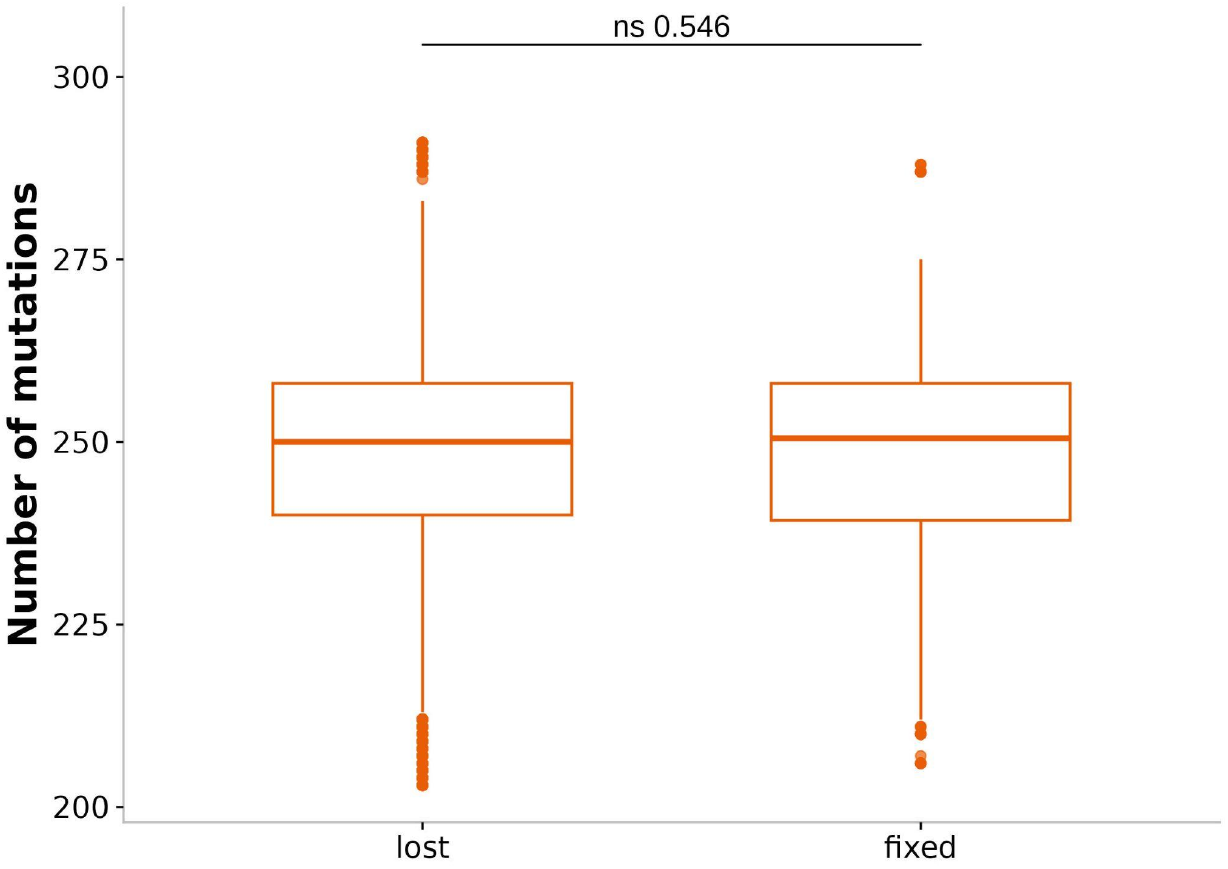
Initial number of partially recessive deleterious mutations in the 1 Mb non-recombining fragments, for fixed vs. non-fixed non-recombining fragments. Results from 1500,000 simulations of the evolution of a panmictic population under a Wright-Fisher model. Each individual is diploid with a 10 Mb chromosome containing a central permanently heterozygous locus. Simulations used a population size (*N*) of 1000, a recombination rate (*r*) of 1 × 10⁻⁹, a mutation rate (*µ*) of 1 × 10⁻⁸, a dominance coefficient (*h*) of 0.3 and a selection coefficient (*s*) of −0.04. The p-value corresponds to a Wilcoxon rank-sum test (ns: non-significant). Boxplots show the median (central line), the 25th and 75th percentiles (box limits), and whiskers extending to 1.5× the interquartile range.

## Supplementary text 1: Notes on Figure 1

Note that the slight excess of heterozygosity, observed even for loci unlinked to the permanently heterozygous locus, and even the neutral ones, likely results from a finite-sample-size effect (Fig 1B). Indeed, observed heterozygosity involves sampling alleles without replacement, whereas expected heterozygosity assumes independent sampling with replacement, leading to a small systematic upward bias in the ratio H_obs_/H_exp_ (J. Wang 2005). More precisely, observed heterozygosity (H_obs_) is calculated from individual genotypes. To determine whether an individual is heterozygous, its two chromosomes are considered sequentially and sampled without replacement. For a biallelic locus with k copies of the focal allele in a diploid population of N individuals, the heterozygosity expected is therefore E(H_obs_) = 2 × [k/(2N)] × [(2N−k)/(2N−1)]. In contrast, expected heterozygosity (H_exp_) is calculated from allele frequency and corresponds to two independent draws: H_exp_ = 2 × p × (1−p) = 2 × [k/(2N)] × [(2N−k)/(2N)]. Consequently, E(H_obs_/H_exp_) = 2N/(2N−1), and E(H_obs_/H_exp_ − 1) = 1/(2N−1). For N = 1000, this corresponds to an expected excess of 0.0005003. This small positive deviation is therefore an inherent finite-population sampling effect of the H_obs_ estimator, rather than a biological signal of excess heterozygosity, and is consistent with the small excess of 0.0005574937 observed at 50 cM. Note also that the small decrease in mean sojourn time at high genetic linkage may be partly attributable to the finite duration of the simulations, as some mutations were still segregating at the end of the 100,000-generation runs (Fig 1A). For example, among mutations with *s* = −0.01 and *h* = 0.1, 5 out of 40,000 were still segregating at 10⁻⁶ cM and 11 out of 40,000 at 10⁻⁵ cM, indicating that the estimated sojourn times are slightly underestimated at these high levels of linkage (orange solid dotted line, Fig 1A). However, it is unlikely to fully account for the observed decrease, which may also partly reflect uncertainty arising from the finite number of simulated mutations.

## Supplementary text 2: Population genetic parameters in natural populations

Estimates of the number of sites under selection in a genome and their recombination rate vary widely depending on the methods used and the species considered (Stapley et al. 2017). For instance, studies in humans suggest that between 8% and 25% of all sites may be non-neutral (Rands et al. 2014; Racimo and Schraiber 2014), whereas studies in Drosophila indicate that more than 50% of sites could be under selection (Halligan and Keightley 2006; Andolfatto 2005). Considering the average chromosome size (∼100 Mb and ∼45 Mb, respectively) in these species, this suggests that the number of sites under selection ranges from ∼8 × 10⁶to 2.5 × 10⁷ per chromosome in humans, and could be above ∼2.3 × 10⁷ per chromosome in Drosophila. Considering the recombination rate (∼1.2 cM/Mb and ∼2.5 cM/Mb) in these species, this corresponds to approximately 6.7 × 10^4^–2.0 × 10^5^ sites under selection per cM in humans and ∼2.0 × 10⁵ sites under selection per cM in Drosophila. However, recombination rates vary by several orders of magnitude along genomes (Halldorsson et al. 2019; Rifkin et al. 2021). For instance, 7% of 100-kb windows in the human genome have recombination rate below 0.01 cM/Mb, which results in a density of selected sites 10⁷ /cM if we consider that 10% of sites are under selection (Halldorsson et al. 2019). Regarding mutation rates, recent estimates across various species are almost all above 10⁻⁹ mutations per base pair per generation, e.g., 7.97 × 10 ⁻⁹ on average in mammals, 4 to 8.4 × 10⁻⁹ in *Drosophila melanogaster* and 1.01 × 10⁻⁸ in birds (Bergeron et al. 2023; Haag-Liautard et al. 2007; Comeron 2014). However, mutation rate variation along the genomes is poorly known and the proportion of deleterious mutations, which matter for pseudo-overdominance establishment, is subject to debate (Conrad et al. 2011; Agrawal and Whitlock 2012; Lesecque et al. 2012; Henn et al. 2015). Nevertheless, most mutations have been estimated to be slightly deleterious, with more than 50% having a selection coefficient *s* between −0.001 and −0.1 (Eyre-Walker et al. 2006; Eyre-Walker and Keightley 2007). For population size, estimates are around 10⁴ for humans (Eyre-Walker et al. 2006; Park 2011), 6 × 10⁵ for mice (Halligan et al. 2010) and 5 × 10⁶for drosophila (Duchen et al. 2013). Dominance coefficients have been estimated to be on average around 0.2, but the distribution of dominance coefficients around this average is unknown (Manna, Martin and Lenormand, 2011; Di and Lohmueller, 2024; Kyriazis and Lohmueller, 2024; see Mrnjavac, Vicoso and Connallon, 2025 for a more detailed discussion).

## Supplementary text 3: Model-based predictions and signature of pseudo-overdominance

The model described here allows us to predict genomic footprints and locations of pseudo-overdominant zones (see also Waller 2021). At the genomic level, pseudo-overdominant zones are expected to display an elevated level of heterozygosity and polymorphism, with numerous deleterious mutations maintained at intermediate frequencies, and strong linkage disequilibrium across the region, with the coexistence of multiple haplotypes within the population. In addition to these genomic features, pseudo-overdominant zones are expected to occur more frequently in regions of high density of sites under selection, low recombination rates and high mutation rates, as well as near loci under balancing selection. Pseudo-overdominance around loci under balancing selection is expected to be more likely in highly selfing species, whereas in other genomic regions it is expected to arise more readily under outcrossing. Pseudo-overdominant zones are also predicted to expand and intensify over time, provided that population genetic parameters are favourable (in particular effective population size, mutation rate and recombination rate). In addition, pseudo-overdominant zones are expected to co-occur with regions of suppressed recombination, such as chromosomal rearrangements, as each can trigger the emergence of the other. Pseudo-overdominant zones are also predicted to result in a reduced fitness of individuals homozygous for a haplotype or recombinant, while heterozygous non-recombinant individuals maintain higher fitness due to the presence of a sheltered load. Altogether, these predictions may help pinpoint pseudo-overdominant zones in genomes and clarify their role in genome evolution.

